# A widespread protein misfolding mechanism is differentially rescued by chaperones based on gene essentiality

**DOI:** 10.1101/2025.09.18.677077

**Authors:** Ian Sitarik, Quyen Vu, Justin Petucci, Paulina Frutos, Hyebin Song, Edward P. O’Brien

## Abstract

Protein misfolding involving changes in non-covalent lasso entanglement (NCLE) status has been proposed based on simulations and biochemical assays of a small number of proteins. Here, we detect hallmarks of these misfolded states across hundreds of proteins by integrating *E. coli* proteome-wide limited-proteolysis mass spectrometry with structural datasets of protein native structures. Proteins containing native NCLEs are twice as likely to misfold, predominantly in regions where these NCLEs naturally occur. Surprisingly, the chaperones DnaK and GroEL do not typically correct this misfolding, except in the case of essential proteins. Statistical analysis links this differential rescue activity to weaker loop-closing contacts in the NCLEs of essential proteins, suggesting misfolding involving these loops is easier to rectify by chaperones. Molecular simulations indicate a mechanism where premature NCLE loop closure, prior to proper placement of the threading segment, leads to persistent misfolded states. This mechanism explains why, in the mass spectrometry data, proteins with NCLEs are more likely to misfold and misfold in NCLE regions. These results suggest widespread NCLE misfolding, that such misfolded states in non-essential proteins can bypass the refolding action of chaperones, and that some protein sequences may have evolved to allow chaperone rescue from this class of misfolding.

## Introduction

Protein folding, the process by which proteins attain their structure and function, has been extensively studied by scientists for half a century^1–4^. Various pathways leading to misfolded states have been documented^5,6^. Therefore, the concept of an unrecognized and potentially widespread class of protein misfolding might seem surprising. However, recent studies^7–10^ utilizing molecular simulations and biochemical assays suggest the existence of a novel class of monomeric protein misfolding involving persistent alterations in non-covalent lasso entanglement (NCLE) structure. Initially proposed in 2022^7^ and supported by indirect evidence from a small number of proteins^7,8,10^, this misfolding mechanism involves the gain or loss of NCLEs within individual proteins. NCLEs consist of a backbone loop closed by one or more non-covalent contacts between residues, with the loop threaded by either the N-terminal or C-terminal segment of the backbone (Fig. 1). According to simulations, NCLEs can misfold in two ways relative to the native structure: either by failing to form a NCLE that is part of the native state or by forming a NCLE that is not native to the structure^8–12^ (Fig. 2c). In both scenarios, long-lived misfolded states may arise as the protein molecule must backtrack^8^ out of these off-pathway states.

**Fig. 1.**
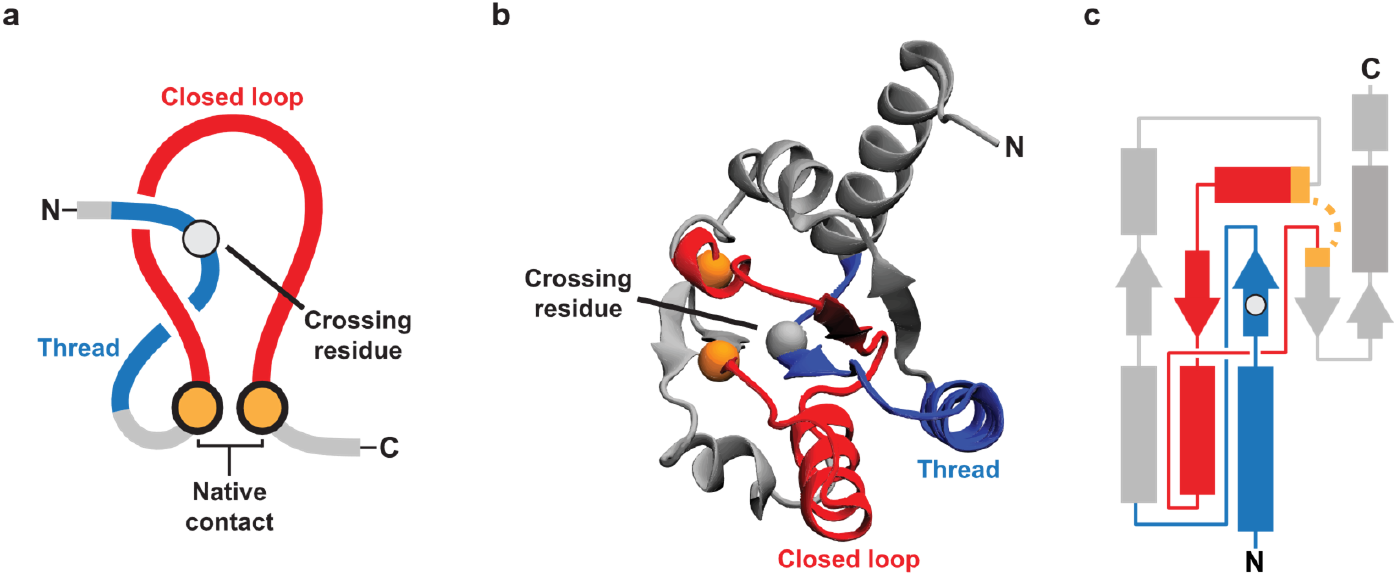
Illustrations of a native NCLE. **a**, The three components of a non-covalent lasso entanglement (NCLE) are (*i*) a loop (red) that is (*ii*) closed by a non-covalent contact (gold), defined by residues with any heavy atoms within 4.5 Ǻ of each other, which is (*iii*) threaded by a segment outside the loop region (blue). The crossing residue is the residue on the threading segment that pierces the plane of the loop. **b**, The crystal structure of the large ribosomal subunit protein uL10 (P0A7J3, PDB 6XZ7, chain H) with the loop (red) closed by residues L72 and A111 (gold spheres – C_*α*_ atom in space fill) that is threaded by a N-terminal segment (blue) with a crossing at V27. **c**, A 2D schematic of the 3D NCLE in b.

**Fig. 2.**
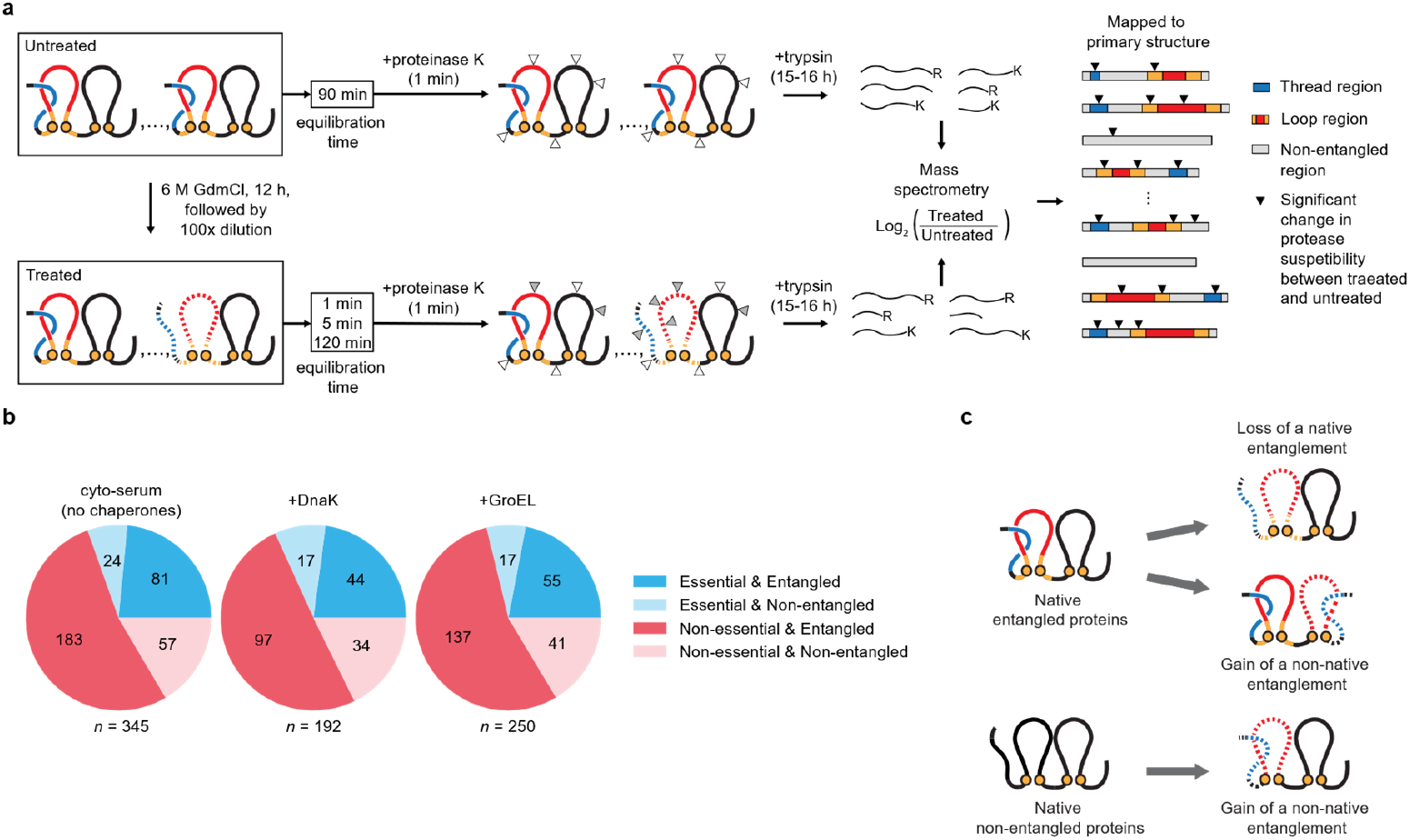
Using high-throughput proteomics data to detect native NCLE misfolding. **a**, Schematic overview of the limited proteolysis mass spectrometry experiments^16^. This method involves splitting a sample of the *E. coli* proteome in two, chemically denaturing one of the samples overnight followed by initiating refolding through a 100-fold dilution. The sample is then allowed to equilibrate for either 1 min, 5 min, or 2 hours, followed by exposure to proteases that cleave surface-exposed residues. The resulting peptide fragments are then identified and their abundances measured via mass spectrometry. This approach detects structural changes within individual proteins at the residue level by comparing differences in the abundances of peptides containing proteinase-K cut-sites observed between the treated sample and the untreated sample. If the patterns are different then the locations of those differences reflect non-native structure at those residues, which we refer to as significant cut-sites. As in Fig. 1, red and blue regions indicate the loop (closed by the native contact shown as orange spheres) and threading regions of a native non-covalent lasso entanglement (NCLE). Dashed lines indicate a non-native NCLE change. White triangles indicate proteinase-K cut-sites that have no statistical difference in their abundance between the untreated and treated samples, while grey triangles exhibit a statistically significant change in abundance. Resulting half-tryptic peptides with significant changes in abundance after false discovery rate correction (black triangles) are mapped to the primary structure of *E. coli* proteins and, in this study, cross-referenced against various NCLE locations and features. **b**, Summary of the native NCLE and essentiality status of the mass-spec observed proteins in the absence of chaperones (cyto-serum) and the presence of DnaK and GroEL. **c**, Possible NCLE misfolding mechanisms of a protein containing a native NCLE, and a protein absent a native NCLE.

High-resolution atomic structures resolved by cryo-electron microscopy (cryo-EM)^13^ would constitute the most compelling evidence of this misfolding class. However, these misfolded states are structurally heterogeneous according to simulations^8,10^, making it difficult for cryo-EM imaging and analysis techniques^14,15^ to resolve them. An alternative approach is to detect the hallmarks of such misfolded states in high-throughput experiments that yield sparse, site-specific structural information.

Here, we utilize data^16^ from high-throughput limited-proteolysis mass spectrometry^16,17^ (LiP-MS), which concurrently probes hundreds of proteins, to investigate protein misfolding at the residue level. This method involves splitting a sample of the *E. coli* proteome in two, chemically denaturing one of the samples overnight followed by initiating refolding through a 100-fold dilution, thereby placing proteins below their denaturant mid-point concentrations. The sample is then allowed to equilibrate for either 1 min, 5 min, or 2 hours, followed by exposure to protease-K that cleaves surface-exposed residues and full digestion with Trypsin. The resulting peptide fragments are then identified and their abundances measured via mass spectrometry^16^ (Fig. 2a). This approach detects structural changes within individual proteins at the residue level by comparing differences in the abundances of (half-tryptic) peptides containing proteinase-K cut-sites^18^ observed between the treated sample and the untreated sample, which presumably consists of mostly folded proteins. If the pattern of cut-sites along a protein’s primary structure is the same between the two samples, then the protein has folded to its native ensemble. If the patterns are different then the locations of those differences reflect non-native structure at those residues^16,17^, which we refer to as significant cut-sites throughout the rest of this study.

Seventy-one percent of globular proteins in *E. coli* possess NCLEs in their folded structure^19^. Moreover, proteins containing native NCLEs have an additional potential misfolding pathway, loss of a NCLE^8^, that proteins without such native NCLEs do not have^8^ (Fig. 2c). Thus, proteins containing native NCLEs have more potential misfolding pathways than those lacking these structural motifs. Based on this, we hypothesize that NCLE misfolding in the *E. coli* proteome will generate more significant cut-sites detected by LiP-MS in proteins with native NCLEs, particularly in regions of the primary structure involved in these NCLEs. Accordingly, we expect to detect statistical associations between the locations of native NCLEs and the sites of non-native structure observed by LiP-MS. This provides a means to identify hallmarks of these misfolded states in a high-throughput manner.

## Results

### Proteins with native NCLE are 200% more likely to misfold

We hypothesized that proteins containing NCLEs in their native state are more likely to misfold compared to proteins that do not. This hypothesis predicts that proteins containing native NCLEs will be more likely to exhibit one or more significant cut-sites upon refolding. To test this prediction, we split the LiP-MS data into proteins that exhibit no significant cut-sites relative to the untreated sample (indicating these proteins refold to their native ensemble), and those proteins that exhibit one-or-more statistically significant cut-sites (consistent with a subpopulation of molecules adopting misfolded conformations). We then computed the odds ratio (OR) (Equation (6.1) in supplemental information) to quantify whether there is a positive statistical association between a protein containing native NCLEs and observed conformational misfolding in the LiP-MS data. We calculate a p-value (*p*) for the significance of this association using the Fisher Exact test^20^. (An OR of 1, with *p* > 0.05, would indicate there is no association, and the presence of a native NCLE has no influence on the presence of significant cut-sites in the sample. A value greater than 1 would mean proteins containing NCLEs are more likely to have significant cut-sites – corresponding to greater misfolding.)

We find an odds ratio of 4.19 (95% confidence interval (CI) = (2.32, 7.62); *p* value that this OR is different than one = 2.93 x 10^-6^, Fisher Exact test, the sample size (*n* = 345) proteins that are both mass spectrometry observable and have a crystal structure) for *E. coli* globular proteins in the absence of chaperones (Fig. 3a). To control for the effects of confounding factors we use logistic regression to determine the odds ratio when accounting for both the amino acid composition of the protein and the protein length, a factor known to positively correlate with longer refolding times^21^. We find an odds ratio of 3.06 (95% CI = (1.55, 6.02); *p* = 0.0012, logistic regression, *n*_prot_ = 345). Thus, proteins that must form NCLEs to reach their native ensemble are 206% more likely to exhibit non-native during the folding process compared to proteins not containing NCLEs of similar size. Because significant cut-sites between the untreated and treated samples arise from structural changes within the protein, we conclude that proteins containing native NCLEs have a greater propensity to misfold compared to proteins not containing NCLEs.

**Fig. 3.**
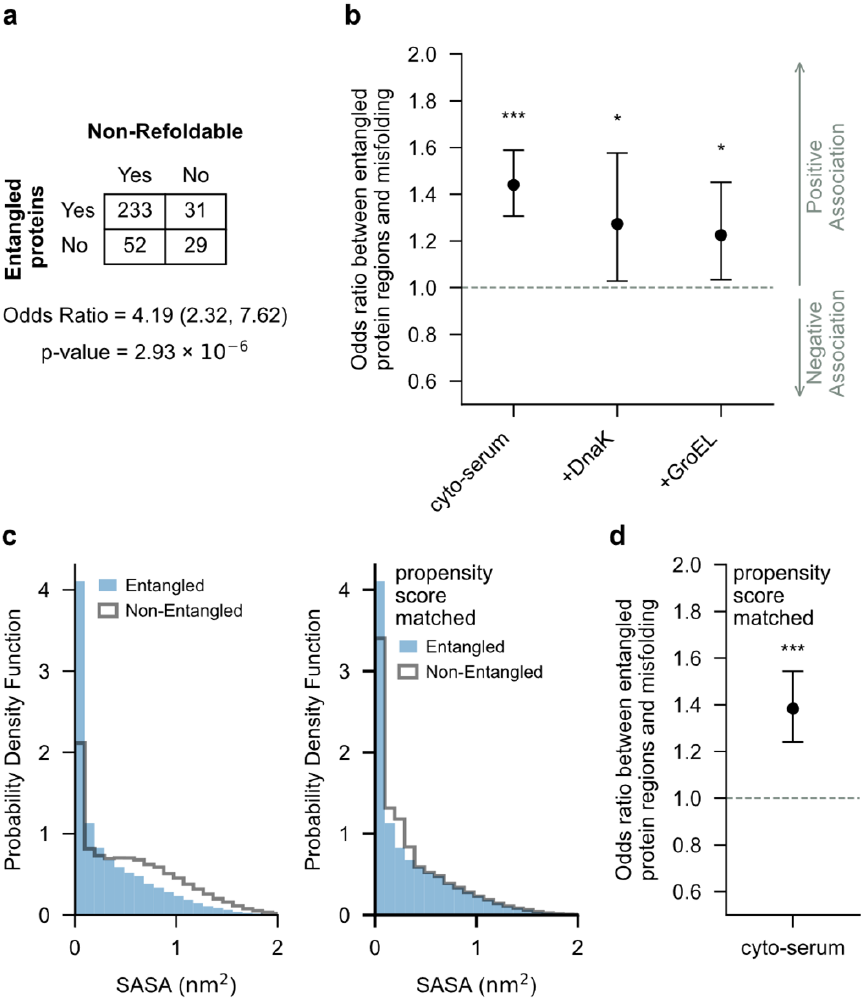
Proteins with native NCLEs misfold more often, biased towards NCLE regions. **a**, Odds ratio (Equation 6.1 in supplemental information) between proteins containing non-covalent lasso entanglements (NCLEs) and misfolding. Shown is the contingency table, odds ratio between the presence of native NCLEs in a protein and that protein being non-refoldable (*i*.*e*., having one or more significant cut-sites), and the *p* value calculated from the Fisher Exact test. **b**, Odds ratio between proteins native NCLE regions and significant cut-sites (indicating misfolding). ***, **, * indicate, respectfully, conditions where the *p* value are below the levels of significance of 0.001, 0.01, 0.05. **c**, Probability density function(s) of the solvent accessible surface area (SASA) for the native NCLE region of the proteome (blue) and the non-NCLE region (transparent with black outline) before Propensity Score Matching (left) and after (right). Propensity Score Matching controls for differences in solvent exposure by constructing a matched dataset of non-NCLE regions with similar SASA as native NCLE regions. **d**, Same as **b** except calculated from the propensity score matched dataset.

### NCLE regions of proteins are 40% more likely to misfold

We hypothesized that the primary structure regions of a protein composing native NCLEs are more likely to misfold than its non-NCLE regions because the NCLE regions have two ways to misfold (loss of the native NCLE or gain of a non-native NCLE) while non-NCLE regions have only one way (gain of a non-native NCLE). This hypothesis predicts that significant cut-sites should occur more frequently in NCLE regions of proteins. To test this prediction, we cross-referenced a database reporting the location of native NCLEs along each protein’s primary structure^19^, derived from crystal structures, against the LiP-MS data (see Methods).

We again use the odds ratio to quantify whether there is a statistical association between the native NCLE regions of proteins and the presence of more significant cut-sites compared to the non-NCLE regions. Here we only consider proteins containing native NCLEs. We find an odds ratio of 1.44 (95% CI = (1.31, 1.59), *p* = 3.10 x 10^-13^, logistic regression, *n*_prot_ = 264 unique proteins in sample with *n*_res_ = 87,716 residues in the model) for *E. coli* globular proteins in the absence of chaperones (Fig. 3b). Thus, the native NCLE regions of proteins are associated with a 44% relative increase in significant cut-sites observed during the folding process compared to non-NCLE regions. Because significant cut-sites arise from structural changes between the treated and untreated samples, we conclude that the NCLE regions of globular proteins exhibit a greater propensity to misfold compared to non-NCLE regions.

### Controlling for residue burial and expression level

Two potential confounding factors could make the previous conclusions erroneous. One is that the NCLE and non-NCLE regions of proteins have different degrees of solvent accessibility in the native ensemble. Threading segments, for example, are surrounded by a loop and hence more likely to be buried in the protein’s native structure compared to non-NCLE regions. These inherent differences could influence the presence of significant cut-sites. To control for this difference, we used propensity score matching^22^, which creates a proteome-wide matched dataset of native NCLE and non-NCLE regions with similar solvent accessible surface areas (Fig. 3c) and calculated the resulting odds ratios (see Methods). With this matched dataset, we find an odds ratio of 1.38 (95% CI = (1.24, 1.54); *p* = 6.79×10^-9^, logistic regression, *n*_prot_ = 264, *n*_res_ = 68,886) in the absence of chaperones (Fig. 3d). Thus, the differences in significant cut-site patterns we previously observed persist in this buffer even after controlling for differences in solvent accessibility. We conclude it is structural changes in NCLEs, not differential solvent accessibility, driving the observed differences upon refolding.

Another potential confounding factor is differences in protein expression levels. Mass spectrometry instruments resolve fewer peptides from low expressed proteins compared to highly expressed proteins. Therefore, low expression proteins can inflate the false negatives in the LiP-MS data and highly expressed proteins can contribute more to the odds ratio. To control for these confounding factors, we restrict ourselves to proteins with at least 50% of the primary structure resolved in the LiP-MS experiments (coverage of the primary structure), and then calculated the odds ratio as a function of the <u>s</u>um of <u>p</u>eptide <u>a</u>bundances (denoted SPA, Equation (5.1) in supplemental information) percentiles. That is, each protein was assigned a score equal to the number of peptide fragments detected and mapped to that protein in the untreated mass spectrometry experiments. That list was then rank ordered, and the odds ratio was computed for proteins at or above a given percentile threshold. We find the odds ratios all remain greater than one and significant in most cases (Supplementary Fig. 4), indicating that, independent of expression level, native NCLE regions of a protein exhibit more significant cut-sites than non-NCLE regions. All results presented in the rest of the study are reported at the 50^th^ percentile SPA for brevity; the robustness of the results to percentile changes are provided in Supplementary Figs. 4 - 11. These controls indicate that the significant cut-sites are arising predominantly from greater misfolding in NCLE regions compared to non-NCLE regions of that protein.

### DnaK/GroEL do not generally correct misfolding bias

Next, we asked if the presence of the chaperones DnaK or GroEL correct the observed bias in misfolding to involve the native NCLE regions in the LiP-MS data. We find odds ratios of 1.27 (95% CI = (1.03,1.58), *p* = 0.027, logistic regression, *n*_prot_ = 141, *n*_res_ = 44,729) and 1.22 (95% CI = (1.03,1.45), *p* = 0.018, logistic regression, *n*_prot_ = 192, *n*_res_ = 62,260), respectively, for *E. coli* globular proteins in the presence of DnaK, and in the presence of GroEL (Fig. 3b, and Supplementary Fig. 4b,c). Thus, even in the presence of these chaperones, NCLE regions still exhibit an association with increased significant cut-sites during the folding process compared to non-NCLE regions. We conclude that DnaK and GroEL do not, across this protein-ensemble average, fully correct the misfolding bias in these NCLE regions.

### Equal misfolding bias in essential and non-essential proteins

In vitro studies demonstrate that DnaK and GroEL help proteins fold^23–25^. Therefore, our ensemble averaged measures must be missing subpopulations of proteins whose folding is catalyzed by these chaperones. We hypothesized that essential proteins are a subpopulation of proteins that are under stronger selection pressure^26,27^ to properly fold and function than non-essential proteins^28,29^, since the functioning of essential proteins are necessary for maintaining life. In contrast, the non-essential proteins can be deleted from the genome with little to no deleterious effects under normal growth conditions^30,31^. Therefore, essential proteins, according to this logic, evolved protein sequences that either (*i*) minimize this class of misfolding or (*ii*) enhance rescue by chaperones relative to non-essential proteins.

To test the first hypothesis we took our dataset and split it into essential^30^ and non-essential proteins and again calculated whether native NCLE regions of essential (or non-essential) proteins were more or less likely to be associated with significant cut-sites upon refolding compared to non-NCLE regions of essential (or non-essential) proteins. We find that in the absence of chaperones, *i*.*e*., in the cyto-serum buffer, both essential and non-essential proteins are equally likely to misfold in NCLE regions compared to non-NCLE regions (Fig. 4), having odds ratios respectively, of 1.45 (95% CI = (1.22, 1.74), *p* = 3.25 x 10^-5^, logistic regression, *n*_prot_ = 81, *n*_res_ = 27,709) and 1.43 (95% CI = (1.27, 1.61), *p* = 2.41 x 10^-9^, logistic regression, *n*_prot_ = 183, *n*_res_ = 60,007). Thus, we reject the first hypothesis, and conclude that in the absence of chaperone quality control mechanisms, essential and non-essential proteins, on average, exhibit misfolding to a similar degree in their native NCLE regions compared to their non-NCLE regions.

**Fig. 4.**
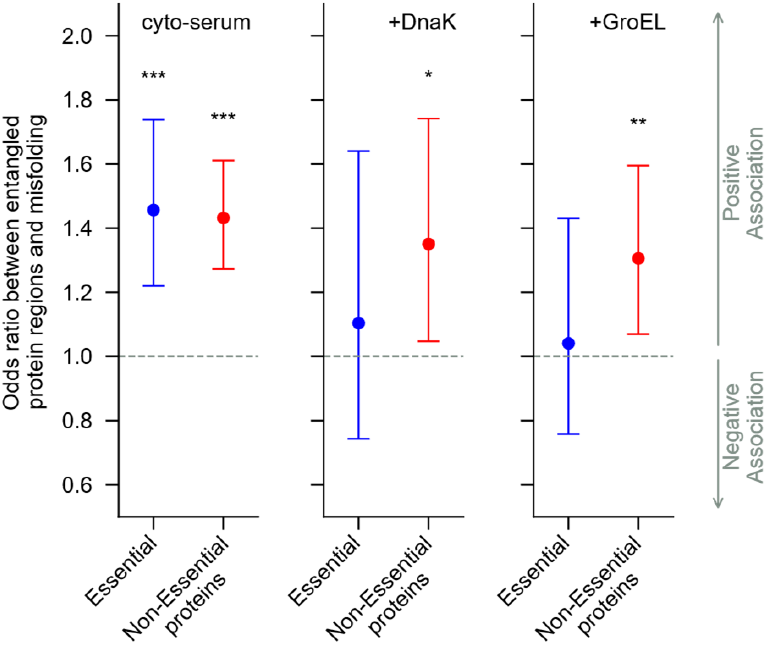
Chaperones correct NCLE misfolding bias in essential but not non-essential proteins. (left) odds ratio between native NCLE regions and presence of significant cut-sites (misfolding) for the set of essential (blue) and non-essential (red) proteins containing native NCLE in the absence of chaperones. ***, **, * indicate, respectively, conditions in which the p-value’s are below significance thresholds of 0.001, 0.01, and 0.05. (middle), same but in the presence of DnaK/J. (right), same but in the presence of GroEL/ES. The raw data used to make these plots were used to calculate the change in the OR results and the significance of the change beyond random chance by permutation testing.

### DnaK/GroEL correct NCLE misfolding bias in essential proteins

We next asked if chaperones correct the NCLE misfolding bias observed in essential and non-essential proteins. To do this, we calculate the change in the odds ratio (ΔOR = OR(Chaperones) - OR(No Chaperones)) going from the buffer in which no chaperones are present to the buffers in which they are present (Fig. 4). The statistical significance of the ΔOR is determined by repeated comparison to randomly permuted versions of the same dataset to compute a one-sided p-value. For essential proteins we find that upon adding GroEL, the odds ratio decreases, with a difference ΔOR = -0.42 (p = 0.028, permutation test). Upon addition of DnaK this difference is ΔOR = -0.35 (p = 0.070). We conclude that GroEL rescues the misfolding bias in native NCLE regions in essential proteins. DnaK exhibits similar behavior, although it is weakly significant, being slightly above our significance level. In contrast, for non-essential proteins the change in odds ratio is much smaller and has larger p-values; when GroEL is present, ΔOR = -0.13 (p = 0.19), and when DnaK is present ΔOR = -0.08 (*p* = 0.31). We conclude that these chaperones do not correct the preferential misfolding in NCLE regions of non-essential proteins.

### Loop-closing residues differ based on protein essentiality

Why is the misfolding bias corrected by chaperones in essential proteins? To address this question we tested four hypotheses rooted in preferential interactions, protein sequence, and protein structure. Our first hypothesis is that essential proteins with NCLEs are more likely to interact with chaperones (often referred to as being ‘clients’ of these chaperones) than non-essential proteins with NCLEs. To test this we used our list of proteins containing NCLEs, cross-referenced it against experimentally identified clients and non-clients of DnaK^31–34^ and GroEL^31,35,36^ (Supplemental Information file 5), and examined if being an essential protein increased the odds of being a client protein. For both chaperones, being an essential protein does not increase the odds of being a client of either as the OR is statistically no different from 1.0 (Fig. 5a). Therefore, essential and non-essential proteins are equally likely to be clients of these chaperones and client status does not explain essential protein chaperone rescue.

**Fig. 5.**
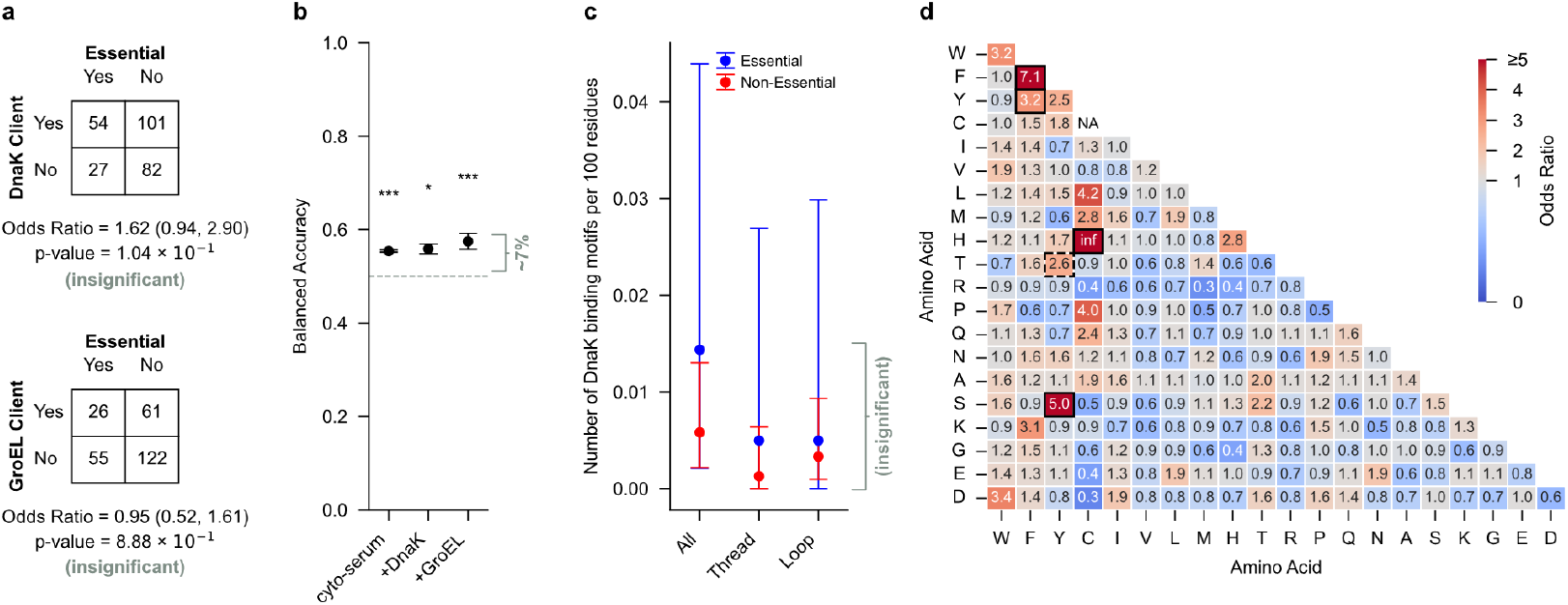
NCLE loop closure and NCLE structure best explain the differential chaperone rescue. **a**, Odds ratio between being a client protein of DnaK (top) or GroEL (bottom) and protein essentiality for the set of proteins that contain native non-covalent lasso entanglements (NCLEs) and are mass-spec observable in the cyto-serum buffer condition. Shown are the corresponding contingency tables, odds ratios, and *p* value from Fisher Exact test. **b**, Least Absolute Shrinkage and Selection Operator (LASSO) balanced accuracy results across 5 k-fold splits of the dataset. 95% confidence intervals are calculated from bootstrapping (10,000 iterations). **c**, Number of Schymkowitz DnaK binding motifs per 100 residues (Equation 4) considering the full-length protein sequence (‘All’), only the native NCLE threads (‘Thread’) or only the native NCLE loops (‘Loop’). Calculated from the set of essential proteins containing native NCLEs (blue) and the set of non-essential proteins containing native NCLEs (red) that were mass-spec observable in the cyto-serum buffer. **d**, Odds ratio (Equation 5) between a particular pair of amino acids, denoted (*i, j*), forming a loop closing contact of a native NCLE and protein essentiality. Those blocks outlined in black have significant *p* values (computed using permutation testing using 100,000 permutations) across all three experimental buffer conditions (*i*.*e*., cytoserum, +DnaK, +GroEL). The block outlined with dashed bolding, (T-Y), had significant *p* values in two out of the three buffer conditions.

Two hypotheses related to sequence could explain the differential rescue. The first is the hypothesis that essential proteins have a higher occurrence of sequence motifs recognized and bound by these chaperones (sometimes called “chaperone-binding motifs”^37^), compared to non-essential proteins. For DnaK we tested this hypothesis by examining if two previously identified DnaK chaperone-binding sequence motifs, listed in Table S2.1, occur more frequently in essential proteins than non-essential proteins. We observed no significant difference (Fig. 5c, Table S2.2). Therefore, DnaK is unlikely to preferentially rescue essential protein misfolding due to an enrichment of DnaK-binding motifs. GroEL has no known sequence-specific client motif, and instead non-specifically binds exposed hydrophobic residues^35,38,39^. This leads to the hypothesis that essential proteins have a larger fraction of hydrophobic residues composing their primary structure than non-essential proteins. We computed the average fraction of hydrophobic residues and found a small, yet significant, difference between the two sets but in the direction opposite of the prediction: essential proteins have 1% less hydrophobic residues than nonessential proteins. It is 62.5% (95% CI = (61.7%, 63.2%), *n*_prot_ = 81) in essential proteins versus 63.6% (95% CI = (63.1%, 64.1%), *n*_prot_ = 183) in non-essential proteins. We therefore conclude that the small differences in the fraction of hydrophobic residues between essential and non-essential proteins do not drive differential GroEL engagement and misfolding rescue of native NCLEs in essential proteins.

The second hypothesis is motivated by the observation^40^ that loop closing contacts of NCLEs are enriched in stabilizing aromatic-aromatic pairwise interactions. We therefore hypothesize that the loop closing contacts in essential and non-essential proteins are different, with non-essential protein enriched in stronger-interacting contacts that could make it more difficult for chaperones to correct their misfolding. To test this hypothesis, we calculated the odds ratio of a given pair of residues (*i, j*) being a loop closing contact in essential proteins relative to non-essential proteins. We find in all three buffer conditions that non-essential-protein-loop-closing contacts are significantly enriched in the stabilizing aromatic-aromatic contacts (F,F), and (F,Y), and aromatic-hydrophobic contacts (S,Y), and (H,C), relative to essential proteins (Fig. 5d). (Also of note, the aromatic-aromatic contact T-Y was enriched in two out of three buffer conditions.) For example, F-F’s odds ratio in the absence of chaperones is 7.14 (*p* = 5.14×10^-14^), meaning there is a 614% increase in the number of (F,F) loop-closing contacts in non-essential proteins relative to essential proteins. Indeed, we find 46.9% (95% CI = (39.8%, 54.5%)) of entangled non-essential proteins contain one or more of these interacting pairs, compared to 19.7% (95% CI = (12.3%, 29.5%)) of entangled essential proteins. Aromatic-aromatic, and aromatic-hydrophobic contacts are known to contribute more to protein free energies of stability than other pairwise classes of interactions, such as polar-polar and polar-hydrophobic interactions^41^. We therefore conclude that while the native, loop closing contacts are stabilizing in both essential and non-essential proteins, the stabilizing interaction energies within these loops are greater in non-essential proteins. This suggests it is energetically easier for chaperones to correct the misfolding in essential proteins than for non-essential proteins.

NCLEs can structurally differ in the number of loop piercing events by the threading segment, loop size, super coiling, as well as many other properties^19^. Each of these structural features has the potential to influence a protein’s misfolding propensity. We therefore hypothesized that the structural nature of NCLEs in essential proteins are distinct from non-essential proteins, and that this could contribute to DnaK and GroEL being able to more easily correct the misfolding in essential proteins. To test this hypothesis, we characterized each native NCLE in our dataset by 18 topological features (Table S2.3) and asked whether those topological features are sufficient to accurately classify a protein as either essential or non-essential. This test is asking whether any of the observed differences between the NCLE properties in these two sets of proteins have sufficient information content to be able to distinguish between essential and non-essential proteins. If they do not have any information content, we conclude they are unlikely to contribute to the observed differential chaperone rescue. We apply the least absolute shrinkage and selection operator (LASSO) regression analysis^42^ on these topological features of NCLEs, and find across the three buffer conditions the regression model has a balanced accuracy ranging from 0.55 to 0.57 (Fig. 5b, Table S2.10). Random classification (i.e., no information content) results in a balanced accuracy of 0.50. Thus, NCLE features provide a 5 to 7% improvement beyond random assignment. We therefore conclude, that while the classification power is small, differences in NCLE properties between essential and non-essential proteins could contribute to differential chaperone rescue.

In summary, the observation that DnaK and GroEL minimize the NCLE-misfolding bias in essential proteins more than non-essential proteins does not arise from differences in their likelihood of being chaperone client proteins or the presence of chaperone-binding motifs. But seems likely to have contributions from less stable loop-closing contacts in essential proteins and differences in the structural nature of these NCLEs.

### Missing crystal structures unlikely to change conclusions

We have restricted our analyses to those *E. coli* proteins that are mass spectrometry observable and have crystal structures available. Globular proteins that have been crystalized have the potential to be systematically different than globular proteins that have not been crystallized. Therefore, our conclusions might only be applicable to this space of already crystalized proteins. To test whether our conclusions are likely to change we expanded our analysis to include proteins that are mass spectrometry observable and have an AlphaFold2 computationally predicted native structure with an average pLDDT score of greater than 70^43,44^. In the case of the buffer absent chaperones, this increases the number of proteins in our dataset from 345 to 554. We find the following conclusions remain robust with this expanded dataset: (*i*) proteins with NCLEs are more likely to misfold than proteins not containing NCLEs; (*ii*) that native NCLE regions of proteins are more likely to misfold than their non-NCLE regions, even after controlling for the potential confounding factors of expression level and residue burial; (*iii*) that DnaK and GroEL do not, on average, correct misfolding of the NCLE regions relative to the non-NCLE regions; (*iv*) that essential and non-essential proteins show equal amounts of native NCLE misfolding; (*v*) that the chaperones rescue the misfolding bias of essential proteins but not non-essential proteins; (*vi*) that there is no association of entangled protein essentiality with being a client protein of GroEL, and (*vii*) that the loop closing contacts in non-essential proteins are enriched in aromatic and hydrophobic contacts relative to essential proteins. We find a single inconsistency with our earlier conclusions. The association of protein essentiality with being a known DnaK client protein is now significant in the AlphaFold2 dataset. The data supporting these claims are provided in supplementary Table 2.4 and 2.5, with results reported at the 50^th^ SPA percentile. Thus, we conclude that nearly all of our conclusions can be extrapolated to those proteins that were mass spectrometry observable in the LiP experiments but do not yet have crystal structures resolved for them.

### Structurally-interdependent loss and gain of NCLEs

What is the misfolding mechanism that explains the structural changes detected in the mass spectrometry data? We have provided statistical evidence that it involves NCLEs, and based on the previous literature, such results are consistent with the hypothesis that the native NCLEs are failing to form. Here, we use the coarse-grained simulation model (Fig. 6a) used in those earlier publications^7–10^ to suggest an answer to this question. As a positive control of the simulation model, i.e., a check that it reproduces known results and thereby assures us of its qualitative accuracy, we simulated two, size-matched sets of proteins. One set corresponding to proteins that were experimentally identified in the LiP-MS data as ‘refoldable’ (*n* = 7), and another set corresponding to similar size proteins that were experimentally identified as ‘non-refoldable’ (*n* = 4). We define refoldable proteins as those lacking any significant cut-sites, while non-refoldable proteins have at least one significant cut-sites. Both sets of proteins have matched pairs of proteins that are within five residues of each other in length, as protein length is a confounding factor known to strongly influence refolding times^21^. We also find that the fraction of misfolded structures (misfolding propensity - Equation (10)) in the non-refoldable set is 63.8% (95% CI = (58.2%, 69.6%)), and in the refoldable set 38.4% (95% CI = (34.1%, 42.8%)) (Fig. 6b). Thus, our model reproduces the experimental observation that the non-refoldable proteins are more likely to misfold than the refoldable proteins. We next asked what is the dominant misfolding mechanism in this model? We do not use the refoldable and non-refoldable protein simulation results, as they are biased to smaller proteins (median size = 135 residues) that are not representative of the typical sizes of proteins in the *E. coli* proteome (median size = 269 residues). Instead, we constructed a third set of proteins that exhibit strong associations of misfolding in native NCLE regions based on the LiP-MS data and randomly selected 10 of these proteins for refolding simulation (see Methods). We analyzed the structures from the last 10% (200 ns) of the quenching trajectories of these proteins to examine the nature of their misfolding. We separate the trajectories into short and long-lived misfolded states based on the median lifetime (Supplementary Fig. 14a) as there is a clear bimodal distribution indicating the misfolded states are either short lived transient states with low structure and high exposure of hydrophobic residues (Supplementary Fig. 14b) or long-lived near-native like states that are likely kinetically trapped. Within the ensemble of long-lived misfolded structures, we observe 26.9% (95% C. I. = (19.8%, 33.4%)) involve solely a loss of NCLE, 9.4% (95% CI = (6.1%, 13.1%)) involve solely a gain of NCLE, and 63.5% (95% CI = (57.0%, 70.6%)) involve both losses and gains of NCLE, i.e., a structure containing both changes of NCLE (Fig. 6e). Thus, a loss of a native NCLE is ∼2.6 times more likely to occur than a gain of a non-native NCLE in the absence of any structural coupling between the two mechanisms.

**Fig. 6.**
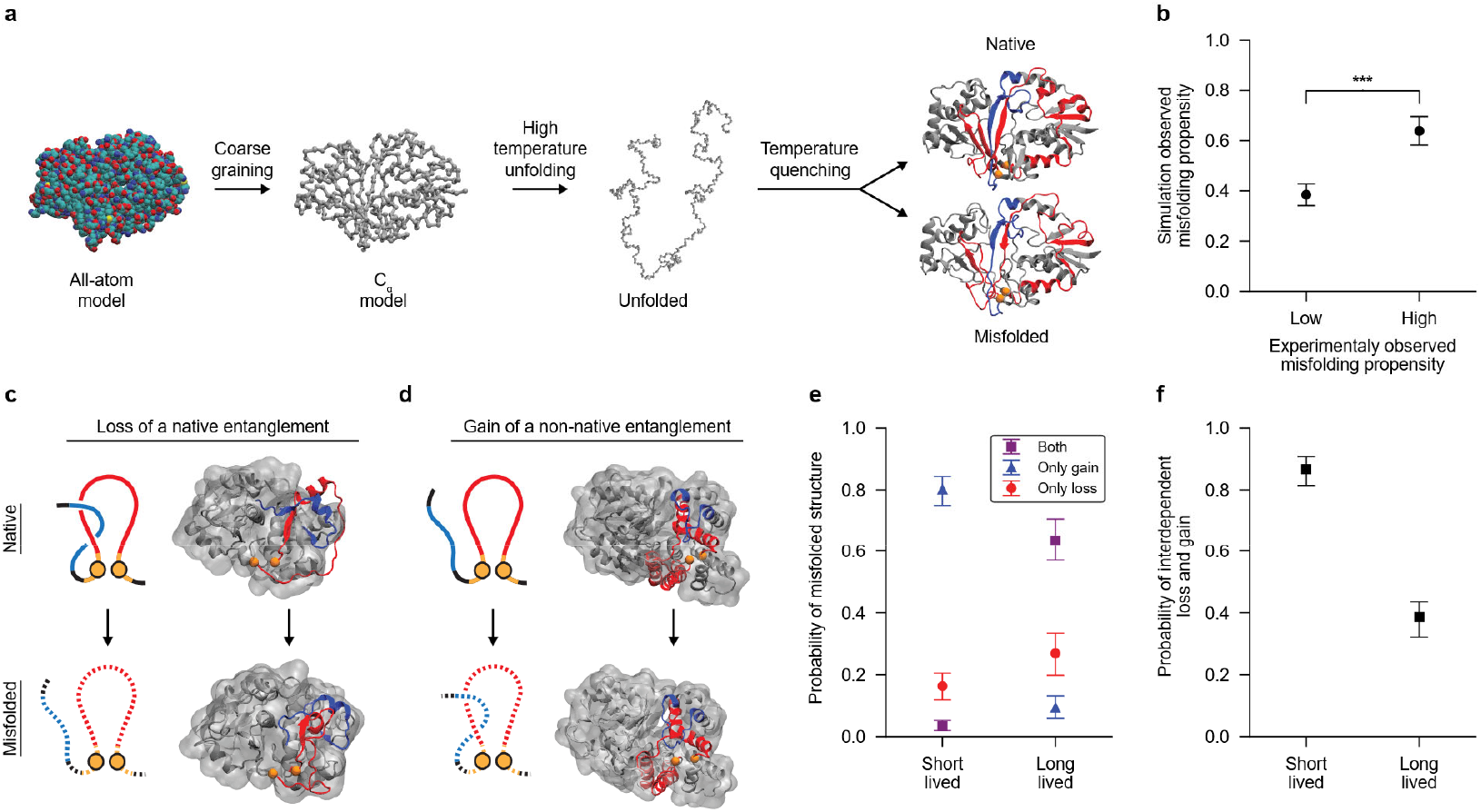
Simulations show dominant misfolding mechanism is structurally-interdependent loss and gain of NCLEs. **a**, Overview of temperature quenching simulations showing the coarse graining (CG) of the all-atom crystal structure using the C_α_ model; unfolding at a temperature above the protein’s melting temperature; then followed by an instantaneous temperature quench to 310 K. Some trajectories properly fold, others adopt non-native misfolded states. Shown is the crystal structure of the protein encoded by gene P37747, (PDB 1I8T, Chain A). Native non-covalent lasso entanglement (NCLE) loop (red) and thread (blue) are shown in the final natively folded and misfolded states to exemplify changes in NCLEs status. **b**, The average misfolding propensity (Equation 10) between the set of proteins predicted to be highly refoldable (low experimentally observed misfolding propensity) and the set of proteins predicted to be highly non-refoldable (high experimentally observed misfolding propensity) (*n* = 50 trajectories for each protein simulated). **c**, Example from the simulations of a loss of a native NCLE with the loop (red) and the thread that is lost upon misfolding (blue) (P31142, PDB 1URH, Chain A). **d**, Example of a gain of a non-native NCLE in the protein from gene Q46856 (PDB 1OJ7, Chain D). The misfolded structure shown is taken from the simulations. **e**, Probability that misfolded structures contain only losses of native NCLE(s), only gains of a non-native NCLE(s), or the presence of both types of misfolding (Equation 11). **f**, The conditional probability of an observed unique loss of a native NCLE having overlap (being structurally interdependent) with a unique gain of a non-native NCLE in the same structure (Equation 12).

**Fig. 7.**
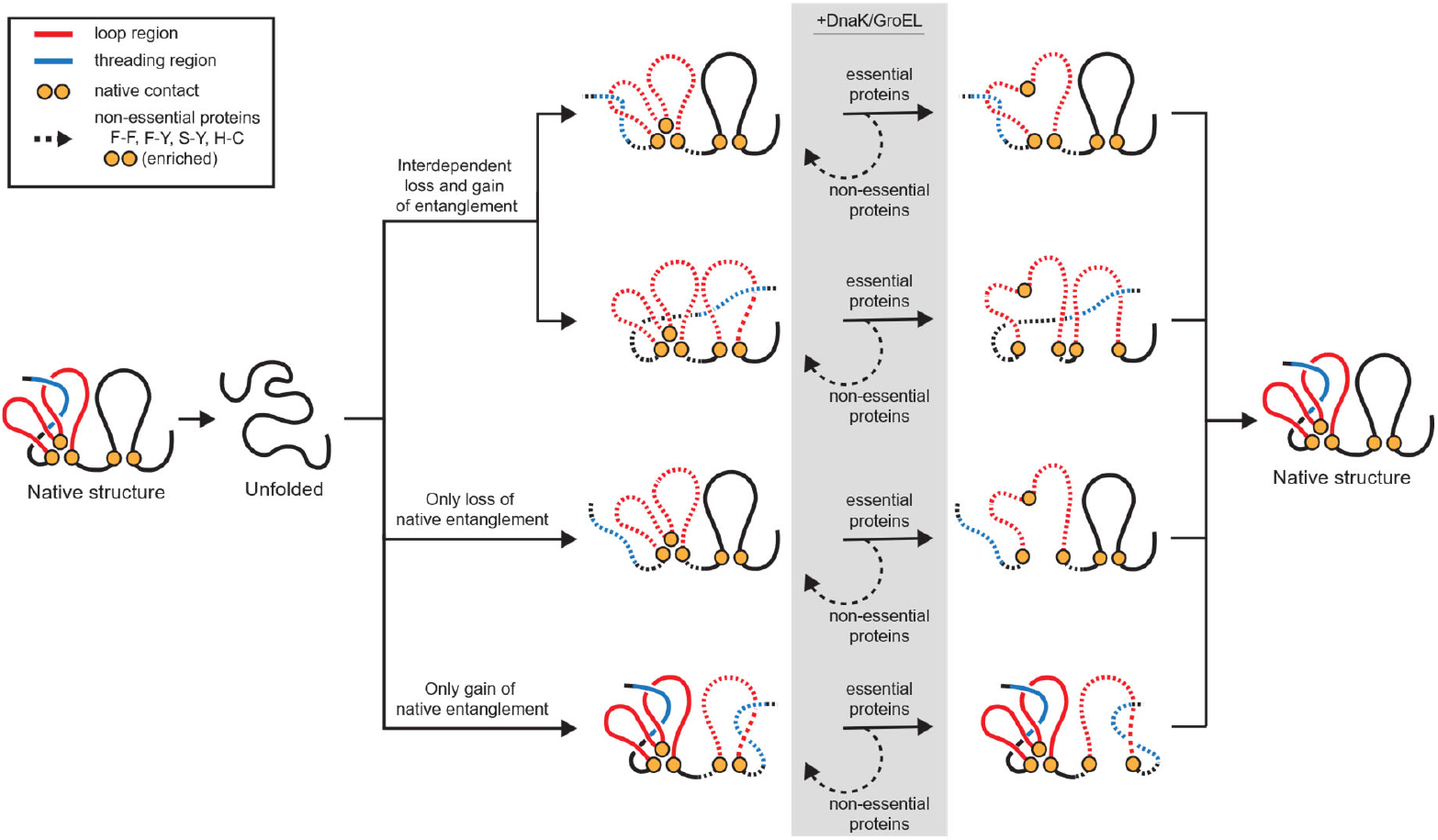
NCLE misfolding model highlights roles of chaperones, essentiality, and loop contacts. Proteins containing native non-covalent lasso entanglement (NCLEs) are more likely to misfold through a mechanism that has a loss of NCLE where the thread or the loop is structurally interdependent with a gain of a non-native NCLE. In the presence of chaperones, the misfolding of native NCLEs in essential proteins is, on average, rescued while it is not in non-essential proteins in part due to the enrichment of aromatic-aromatic and hydrophobic-aromatic loop forming contacts (F-F, F-Y, S-Y, H-C). We speculate this is due to chaperones DnaK and GroEL having to expend less energy in reopening the loop of essential proteins than non-essential proteins to allow for proper threading of the native NCLE. This diagram only displays misfolding pathways. There are, of course, other pathways involving on-pathway intermediates that are not shown in this model.

We examined if losses and gains of NCLE are dependent on each other or occur independently along the primary structure. To assess this, we calculate the conditional probability that, given both a loss and gain of NCLE has occurred in a structure, that one or more segments that compose the lost native NCLE directly take part in the new gain of non-native NCLE. If independent, this probability will be zero. We find that this probability is 61.4% (95% CI = (57.1%, 67.2%)) and a probability of a loss happening in isolation is 38.6 (95% CI = (32.2%, 43.5%)) (Fig. 6f). We conclude that the dominate misfolding mechanism in this model is loss of a native NCLE, and two thirds of the time this loss is directly involved in the simultaneous gain of a non-native NCLE. This is consistent with and provides an explanation for the LiP-MS observations that misfolding occurs more frequently in proteins containing native NCLEs and in the NCLE regions.

## Discussion

This is the first study to detect the hallmarks of this class of entanglement misfolding in a high-throughput manner. LiP-MS yields sparse, site-specific structural information. However, its ability to simultaneously probe many proteins for non-native structural changes means the large data sets it produces are amenable to statistical analyses that can relate misfolding propensities to the structural properties of the native state^16,17^. This approach has allowed us to make two discoveries. The first is that proteins containing native NCLEs are far more likely to misfold. The second is that misfolding in these proteins predominantly is localized to the NCLE regions along the primary structure. These two results are significant because they provide experimental evidence that misfolding involving changes of non-covalent lasso entanglement status could be a widespread phenomenon.

A predominant paradigm in molecular biology is that when a protein misfolds it will either aggregate^45,46^, get degraded^47,48^, or chaperones will help it fold^23–25,32^. This paradigm has been suggested to be incomplete based on a meta-analysis of the experimental literature^9^. Twenty *in vitro* studies examining the impact of chaperones on folding have found, consistent with this paradigm, that some proportion of the molecules are aided in their folding process. However, inconsistent with this paradigm, these studies also found there is always a subpopulation that is not aided, remaining soluble, non-functional and hence misfolded. In this study, we were able to test whether the *E. coli* chaperones GroEL/ES and DnaK/J correct the bias in misfolding in native NCLE regions and found they do not at the proteome-wide level. Surprisingly, for essential proteins these chaperones do tend to correct their bias in misfolding, but not so for non-essential proteins. This difference does not arise from essential proteins being a preferred client of these chaperones, nor from essential proteins having more chaperone binding motifs within their protein sequence. Instead, our results indicate that one contributing factor is the identity of the pairs of interacting residues closing the loop of the native NCLE. Non-essential proteins are more likely to have aromatic-aromatic and aromatic-hydrophobic contacts closing the loop compared to essential proteins (46.9% versus 19.7%). Such contacts are, in relative terms, highly stabilizing^41^. This suggests that one reason these chaperones can rescue the misfolding in essential proteins is because they can expend less free energy to open up a prematurely closed loop providing another opportunity for the threading segment to attain its proper native positioning. Conversely, this process is less efficient for non-essential proteins that contain these strongly stabilizing contacts. Indeed, the notion that energy must be expended to unfolded misfolded conformations is one of the two key mechanisms of GroEL’s refolding action^49–51^. This suggests the hypothesis that the primary structure of essential proteins may have evolved to be less stable, not to avoid NCLE misfolding (we found NCLE misfolding has similar odds ratios in essential and non-essential proteins in the absence of chaperones), but to allow the refolding action of chaperones to efficiently correct such misfolding.

Another factor that might contribute is differences in native state NCLE structural properties (Table S2.10). These differences were, in terms of classification power, weak – yielding just a 5 to 7% increase above random classification (Fig. 5b). The relative changes across the topological features do not lend themselves to an obvious molecular hypothesis of how they could contribute to differential chaperone rescue. This suggests the possibility that these differences may not be a causal driving force. In future studies, targeted biophysical experiments could be useful in establishing a mechanism for the observed differential rescue of essential proteins.

This study was motivated by our hypothesis that there are more ways to misfold a natively entangled protein than a protein whose native state does not contain any native NCLEs, and that this should manifest itself in the LiP-MS data as a statistical bias towards more frequent structural changes being observed in proteins containing native NCLEs and NCLE regions. Specifically, we predicted that a failure-to-form mechanism would be the dominant mechanism. While our simulation results are broadly consistent with this prediction, the simulations suggest a more complicated molecular scenario is giving rise to the LiP-MS signals involving correlated misfolding events. Specifically, while 26.9% of misfolded structures involve solely a loss of a native NCLE, 63.5% of the remaining misfolding events involve the structurally-interdependent loss of a native NCLE and gain of a non-native NCLE involving the thread of the native NCLE that failed to form. To illustrate, consider the opposite case: structurally independent loss and gain of NCLEs. In this case, along the primary structure of a protein, one segment fails to form its native NCLE, and a non-overlapping segment of the sequence gains a non-native NCLE. These occur structurally independently. What we observe in 61.4% of the cases is that such simultaneous loss and gain of NCLE involves overlapping backbone segments. Specifically, the thread of the native NCLE failing to position itself within the native loop, and instead structurally taking part in the newly formed non-native NCLE. Thus, in this way, even gains of non-native NCLEs, can still be structurally biased towards native NCLE regions, and contribute to the increased odds observed in the LiP-MS data that non-native structural changes will be associated with native NCLE regions.

In statistical association studies, such as this, there are common misinterpretations that can arise. Our results do not mean that all proteins containing NCLEs misfold, and that if they do misfold they all misfold involving a failure to form mechanism. Our results show that there is a bias - if these structural elements are present in the native state, they are more likely to misfold compared to proteins or protein backbone segments that do not have them. Misfolding involving gains of NCLE outside the native NCLE region can still be present, albeit to a relatively lesser extent. Further, while the presence of DnaK/J and GroEL/ES eliminate the association between misfolding and native NCLE regions in essential proteins, this does not mean they eliminate all misfolding in these proteins. Rather, we find misfolding is equally likely in these two regions in the presence of these chaperones. And finally, while we have provided evidence that changes of NCLE status contribute to the misfolding in the set of proteins in the data, this does not preclude other misfolding mechanisms from simultaneously being present in these systems, such as those caused by proline isomerization, incorrect disulfide bond formation, etc.

The present study does not account for co-translational folding and the influence of the ribosome on protein folding. Previous simulation studies comparing NCLE folding on and off the ribosome found that a majority of the misfolded state populated during refolding are similar to the misfolded states populated co- and post-translationally^7–9,12^. This suggests that the conclusions of this study might be applicable in the context of co- and post-translational folding and chaperone interactions. However, it is important that this speculation be tested in a future study.

This study is unable to quantify how widespread this misfolding bias is across the proteome. To quantify this, we need to know at the individual protein level whether it exhibits a misfolding bias in its native NCLE regions compared to its non-NCLE regions. However, the sparsity of signal (i.e., the number of significant cut-sites exhibiting a statistically significant change between treated and untreated samples) at the individual protein level does not provide sufficient statistical power to detect such biases. For example, in the cyto-serum dataset, there are a median of 3 cut-sites per protein. It is only when the data is pooled across proteins that the statistical associations can be detected. This is a common limitation in high-throughput studies with sparse signals^52,53^. We expect, however, that this misfolding mechanism is present in a majority of globular proteins. We base this on several observations. First, all twenty studies in our previous meta-analysis^9^ detected subpopulations of kinetically trapped, soluble misfolded proteins that were not acted upon by chaperones. Second, synonymous mutations have been repeatedly observed to alter the fraction of soluble protein molecules that are active^8,54,55^. Third, high-throughput protein folding simulations estimate the majority have subpopulations that become misfolded and kinetically trapped^7–10^. These three observations indicate that soluble, kinetically trapped misfolded subpopulations are widespread. Changes of NCLE status provide a simple unifying explanation for all of these observations.

Such widespread misfolding may be relevant to protein homeostasis, disease, and aging. Cellular proteostasis processes are likely to respond to grossly misfolded structures. Therefore, we expect that proteins with native NCLEs are more likely to be degraded than those without. Indeed, this has been observed in yeast under heat shock in which such nascent proteins are at 200% greater odds of being degraded than similar proteins that do not need to form native NCLEs^56^. But even in the absence of such cellular stress, we predict they will be more likely to be degraded due to their increased propensity to misfold. Similarly, this perspective also predicts that proteins that aggregate are more likely to be enriched with proteins that misfold through a failure-to-form mechanism. In contrast, near-native misfolded structures – which resemble the native structure but have structurally localized changes in NCLE - are more likely to remain soluble and bypass these proteostasis processes^7–9^. As mentioned, this more subtle form of misfolding has been proposed as an explanation for how misfolded states can bypass the refolding action of chaperones, and the long-term changes in enzyme structure and activity when encoded by synonymous-mRNA variants. More broadly, the loss of function that can occur with such misfolding has the potential to contribute to disease and aging. Previously discovered misfolding mechanisms, in which the gene product is no longer as efficient in its activity, have been found to contribute to various diseases^45,57,58^, and there is no reason to think an NCLE mechanism of misfolding cannot have similar outcomes. Aging has multiple molecular and cellular hallmarks, one of which is age related structural changes in some proteins^59–61^. The nature of these structural changes is unclear. We hypothesize that they may involve changes in non-covalent lasso entanglement status. Thus, the biological consequences of this class of misfolding could be wide-ranging.

## Methods

### Determining NCLEs across 1294 representative proteins

We take a set of 1,294 high quality (resolution ≤ 3 Å) protein structures from the Protein Data Bank (PDB) that were selected to have at least one alignment with the UniProt^62^ canonical sequence with a minimum of 95% identity and positive scores, less than 5% gaps, and an Expect score of less than or equal to 10^-5^. We remove structures that are identified as membrane or transmembrane by Uniprot and remove structures with more than 50% sequence disorder as determined by MobiDB^63^. We identify all native NCLEs present by the Gauss Linking Integration method^64^. The loop of an NCLE is defined as those residues between residues *i* and *j* that are in contact with each other. Two residues are in contact if they have any heavy atoms within 4.5 Å of each other.

The N and C terminal segments flanking this loop are then examined for any linking with the loop with linking values denoted as *g*_N_(*i, j*) and *g*_C_(*i, j*). We calculate these values using the partial Gauss double integration method created by Baiesi and co-workers^40,65^. In brief, for a given structure of an *N*-length protein, with a native contact present at residues (*i, j*), the coordinates *R*_*l*_ and the gradient *dR*_*l*_ of the point *l* on the curves were calculated as:

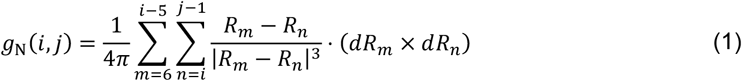

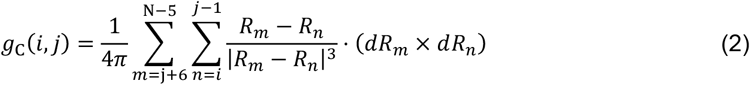

A given loop is determined to be entangled if |*g*_N_(*i, j*)| ≥ 0.6 or |*g*_C_(*i, j*)| ≥ 0.6. If an NCLE is detected for a given native contact, we use the Topoly^66^ Python package to identify the residues at which the termini cross the loop plane. We obtain unique NCLEs using a previously published clustering method^19^ (Supplemental methods 1.3). We filter AlphaFold structures for both high quality structures and NCLEs (Supplemental methods 1.2).

### Determining significant cut-sites across the proteome

To determine which sites across the proteome show a significant change in proteolysis susceptibility we examine the high throughput limited proteolysis mass spectrometry (LiP-MS) dataset previously published by the Fried Lab^16^. The proteome discover (PD) data was reanalyzed using the FLiPPR^18^ protocol to determine unique proteinase-K cut-sites that have significant changes in abundance between an untreated native sample and a sample refolded through treatment with 6 M Guanidinium chloride and dilution jump into cytosol like medium. In brief, a significant cut-site must have a significant increase or decrease greater than or equal to a 2-fold change in the abundance between the untreated sample and a treated sample in at-least 1 of the refolding timepoints (1 min, 5 min, 2 h) after FDR correction using the Benjamini-Hochberg method^67^. This pooling of the timeseries data increases the statistical power to detect associations between native entangled structure and adoption of non-native structure in the experiment. Even in a limiting case in which the non-native ensemble is different at each time point, underlying associations between NCLEs and misfolding can still be detected. At least 72% of the proteins exhibiting non-native structure (Supplementary Fig. 2) and 40% of their significant cut sites (Supplementary Fig. 3) are common across these timepoints.

### Controlling for changes in protein abundance

Controlling effects of endogenous protein abundance differences in the LiP-MS data is done through both the sum of the peptide ion abundances (SPA) and the peptide coverage determined by proteome discover (PD). For most analysis in this work we control for changes in natural protein abundances by rank ordering the proteins based on their SPA and sweep every 10^th^ percentile taking subsets of proteins with abundances greater than or equal to the percentile in a given buffer condition are considered in downstream analysis. Furthermore, we only consider proteins that have at least 50% of the canonical protein sequence correctly identified by PD to help mitigate false negatives. Finally, for all work reported in the main text we only report the results from the median SPA threshold and the sweeps of the full threshold can be found in the supporting information. We note that this choice of thresholds to control for protein abundance does not bias the protein length or number of domains in our dataset.

### Modeling observed significant cut-sites across the proteome

We fit a simple logistic regression model using the statsmodel^68^ python package to estimate the log-odds of observing a significant cut-site as a function of the amino acid type and whether the residue was in the native NCLE region of the proteome or not.

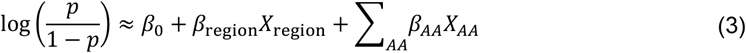

Where *X*_region_ is a binary variable defining if the residue was in the native NCLE region of the protein or not, and *X*_*AA*_ is a binary variable defining if the residue was of the type *AA* where *AA* is one of the 20 canonical amino acids.

An odds ratio is then defined as, 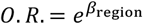 where *β*_region_ is the resulting coefficient of the regression variable describing which region a given residue belonged in the binomial regression. (See supplemental information for more details.)

### Essential and non-essential protein definition

We define our set of essential proteins as those identified in the knock-out experiments of Baba T and collaborators^30^. They identified 296 essential proteins of which 188 are present in our set of high-quality crystal structures. All other proteins are defined as non-essential.

### Scanning for DnaK binding motifs

For each protein we created three sets of sequences to search for potential DnaK binding motifs: (1) the whole FASTA sequence of the protein, (2) the sequence composing the loop of each unique NCLE, and (3) the sequence composing the threading segments of each unique NCLE, where the sequence for each thread was taken as the crossings that pierced the loop plane (as determined by Topoly) +/- 10 residues. We scanned each set of sequences for matches to three different motif patterns previously identified from a combination of experiments^37^, homology modeling^69^, and an expanded Bukau-like motif that follow the pattern BHHHHHB, where B are basic residues and H are hydrophobic residues (Table S2.1, Supplemental methods 1.10).

To compare the sets of essential and non-essential proteins we calculate the number of motifs per 100 residues as

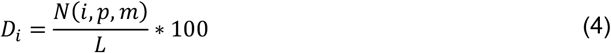

Where *i* is the index of the protein, *p* is the pool of sequences (i.e. whole sequence, just the loops, or just the threads), *m* is the motif, *N*(*i, p, m*) is then the number of motifs in that pool for that protein, and *L* is the length of the protein. Confidence intervals for these distributions for any given set of proteins were calculated by basic bootstrapping (*n* = 10,000) and the test of a difference between the two distributions was done by permutation (*n* = 10,000). Finally, we corrected the permutation p-value’s by the benjamini-hochberg^67^ procedure to control for false discovery rates.

### Modelling protein essentiality with NCLE features

For each protein with a native NCLE, we cluster the set of ‘raw’ native NCLEs into sets that represent the same topological NCLE and choose one with a minimal loop as the representative native NCLE for each set^19^. For each of these unique NCLEs we calculate 18 structural features designed to capture the complexity of a native NCLE, (Table S2.3).

To determine which, if any, features best discriminate between the classes of essential and non-essential proteins we use the Least Absolute Shrinkage and Selection Operator (LASSO) logistic regression model to find an optimal set of parameters. We scanned the inverse regularization strength *C* from 0.00001 to 10 in 1,000 evenly placed steps. To determine if the average balanced accuracy across 5 stratified k-folds of the standard scaled data for a given regularization strength is significant beyond random chance we perform the LASSO regression method described above on randomly permuted versions of the dataset (*n* = 10,000) and determine what is the probability of observing an average balanced accuracy that is as different from 0.5 by random chance. The optimal values of *C* that provided the maximal balanced accuracy with the minimal number of non-zero robust features in the absence of chaperones was 1.5, in the presence of DnaK 2.5, and the presence of GroEL 2.5.

### Loop closing contact hydropathy and contact enrichment

For any two pairs of amino acids (*a, b*) that form loop closing contacts in either the set of essential or non-essential proteins used in this study we determine if there is any statistically significant enrichment in either set by calculating the odds ratio for the association of a given pair of amino acids (*a, b*) being a loop closing contact and the protein being essential.

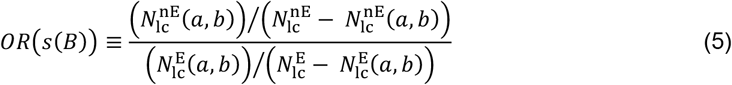

Where *s*(*B*) is the set of proteins that have a SPA greater than the 50^th^ SPA percentile and have more than 50% of their primary sequence observed in the LiP-MS experiments in buffer condition 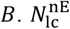 and 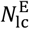 are the number of loop-closing contacts in the set of non-essential and essential proteins respectfully. 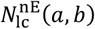and 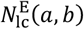is the number of loop-closing contacts composed of residue types (*a, b*) in the set of non-essential and essential proteins respectfully.

We estimate a p-value by permutation of the loop-closing contacts (n = 100,000) and apply a false discovery rate correction using the benjamini-hochberg procedure. We consider the pairs that are significantly enriched or depleted to have an adjusted p-value less than 0.05 in all three buffer conditions (cyto-serum only, +DnaK, and +GroEL) to ensure we are only considering the enrichment of the most robust pairs of loop closing amino acids observed across all experiments.

### Calculating fraction of native contacts (*Q*)

The fraction of native contacts (*Q*) is calculated as a ratio of native contacts present in a given structure relative to a reference “native” structure:

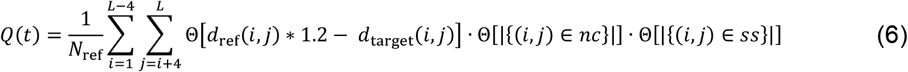

Where *i* and *j* are residue indexes and *L* is the length of the protein. The set of native contacts in the reference structure *nc* is defined by those pairs that have a Euclidean distance between their alpha carbons *d*_ref_(*i, j*) ≤ 8 Å, at least 3 residues apart along the primary structure, and be located within the set of secondary structures residues *ss* defined by STRIDE^70^. For a given reference structure there are *N*_ref_ native contacts made. Θ is the Heaviside step function.

### Calculating fraction of native NCLE status changes (*G*)

The fraction of native contacts (*G*) that have a change in one of the terminus entanglement status is calculated as:

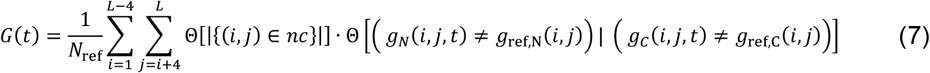

Where *i* and *j* are residue indexes and *L* is the length of the protein. The set of native contacts in the reference structure *nc* is defined by those pairs that have a Euclidean distance between their alpha carbons *d*_ref_(*i, j*) ≥ 8 Å and at least 3 residues apart along the primary structure. The number of native contacts is in the reference structure is *N*_ref_. The partial linking values for the N-terminus and C-terminus are *g*_*N*_ or *g*_*C*_ respectively. Θ is the Heaviside step function.

### Calculating misfolding propensity

The misfolding propensity is calculated for the last 200 ns of each quenching trajectory *t* of protein *i* as the fraction of non-native frames identified by the simple thresholding of two key yet effective order parameters, the fraction of native contacts (*Q*) and the fraction of native contacts with a change in NCLE (*G*). We define our thresholds on 10 independent reference simulations at 310 K in the following manner:

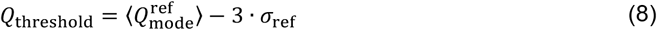

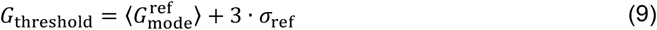

Where 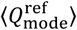 and 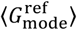 are calculated by taking a sliding window of size 15 ns and obtaining the mode of each order parameter across each reference simulation and then computing the average of this series of modes and standard deviation *σ*_ref_. We then select structures with *Q* ≥ *Q*_threshold_ and *G* ≤ *G*_threshold_ (supplementary tables 2.7 - 2.8) and designate them as native structures while everything else is considered non-native. This approach is consistent with our previous publications^7,9^. The misfolding propensity is then:

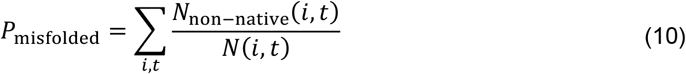

Where *N*_non-native_(*i, t*) is the number of non-native structures and *N*(*i, t*) is the number of frames in the last 200 ns of the quenching trajectory.

We assess the significance of the difference in the misfolding propensity between candidate sets (I) and (II) (defined in the “Selecting candidates for coarse grained molecular dynamics simulations” section) by permutation (*n* = 10,000) and obtain 95% confidence intervals via basic bootstrapping.

### Calculating probability of misfolding mechanism type

To calculate the probability of observing either a loss of a native NCLE or a gain of a non-native NCLE we first filter the trajectories remove those that are natively folded in the last 200ns of trajectory *t* of a protein *i* in candidate set (III) (Supplemental methods 1.13). Trajectories are considered natively folded and discarded if 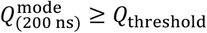 and 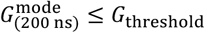, where *Q*_threshold_ and *G*_threshold_ thresholds are set by the reference simulations (Equations (8) and (9), supplementary tables 2.7 - 2.8). For those trajectories that pass the filtering criteria we calculate the average lifetime of the misfolded state in each trajectories as the average fraction of consecutive frames where a change of NCLE status was identified multiplied by 0.075 ns as the timestep. We then separate the trajectories into long lived and short-lived misfolded states based on the median lifetime of ∼100 ns.

To calculate the probability of a given structure exhibiting a misfolding mechanism *M* (only a loss of a native NCLE, only a gain of a non-native NCLE, or both) within each trajectory *t* we use the following:

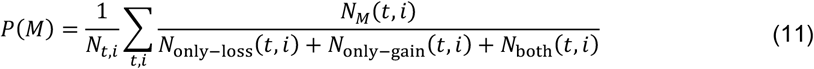

Where *N*_*M*_(*t, i*) is the number of structures observed in the quenching trajectory *t* of protein *i* with the particular misfolding type *M* present and *N*_*t,i*_ is the total number of trajectories across all protein in a given dataset considered. We then report the ensemble average of *P*(*M*) across all proteins and quenching trajectories that met the criteria above.

Considering only the structures that contain both a loss of a native NCLE and the gain of a non-native NCLE we calculate the conditional probability that a loss is structurally interdependent with a gain in a given trajectory *t* by examining whether the key topological features (i.e. the loop closing native contacts +/- 3 residues and the crossing residues +/- 3 residues) of the NCLEs are shared. The conditional probability of a loss of a native NCLE being interdependent with at least 1 gain of a non-native NCLE in the last 200 ns of trajectory *t* of protein *i* is then given the structure had both present is then defined as:

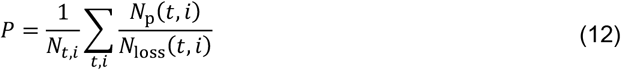

Where *N*_p_(*t, i*) is the number of losses of NCLE that had primary structure overlap of key NCLE topological features with at least 1 gain of NCLE in the same structure, *N*_loss_(*t, i*) is the total number of losses observed across all structures containing both a types of misfolding, and *N*_t,i_ is the total number of trajectories across all protein in a given dataset considered.

## Data Availability statement

The datasets used in the statistical analysis are provided as supplementary files 1-6 and on the associated GitHub repository (https://github.com/obrien-lab-psu/Failure-to-Form_Native_Entanglements ). Molecular dynamics trajectories are available upon request as they are to large to store publicly. The mass spectrometry data analyzed in this paper was taken from the PRIDE repository PXD030869.

## Code Availability statement

The code is shared on GitHub (https://github.com/obrien-lab-psu/Failure-to-Form_Native_Entanglements ).

## Acknowledgements

E.O. gratefully acknowledges support from the National Science Foundation (MCB-2031584) as well as from the National Institutes of Health (R35-GM124818). Portions of numerical computations and data analysis in this work have been carried out on high-performance computing collectively known as ROAR, which is operated by the Institute for Computational and Data Sciences at The Pennsylvania State University.

## Credit authorship contribution statement

**I.S**.: Investigation, Methodology, Formal analysis, Visualization, Validation, Data curation, Writing – original draft, Writing – review and editing.

**Q. V**.: Methodology, Writing – review and editing.

**J. P**.: Methodology, Writing – review and editing.

**P. F**.: Formal analysis, Methodology

**H. S**.: Methodology, Validation, Writing – review and editing

**E. O**.: Conceptualization, Formal analysis, Funding acquisition, Investigation, Methodology, Project administration, Resources, Supervision, Validation, Writing – original draft, Writing – review and editing.

