## Supplemental Information for "A widespread protein misfolding mechanism is differentially rescued by chaperones based on gene essentiality"

### 1. Supplemental Methods

#### 1.1. Determining unique non-covalent lasso entanglements across the set of 1294 representative protein structures

We take a set of 1294 representative protein structures from the PDB and first calculate all native NCLEs present by the Gauss Linking Integration method[1]. Given a protein of length $N$ a native NCLE is defined by two topological features, a loop segment and a threading segment that pierces the plane of the loop. A loop in a protein structure is defined as being composed of the backbone trace connecting residues *i* and *j*, which have formed a native contact in a given protein conformation. The native contact between *i* and *j* is considered to close this loop, even though there is no covalent bond between these two residues. We define a native contact as any two residues with heavy atoms within 4.5Å of each other.

Outside this loop is an N-terminal segment, composed of residues 1 through *I* - 1, and a C-terminal segment composed of residues *j* + 1 through $N$. These two segments represent open curves, whose entanglement through the closed loop we characterize with linking values denoted as $g_{\mathrm{N}}(i,j)$ and $g_{\mathrm{C}}(i,j)$. We calculate these values using the partial Gauss double integration method proposed by Baiesi and co-workers[2,3].

For a given structure of an *N*-length protein, with a native contact present at residues $(i,j)$, the coordinates $R_l$ and the gradient $dR_l$ of the point $l$ on the curves were calculated as:

$$R_l = \frac{1}{2}(r_l + r_{l+1}) \tag{1.1}$$

$$dR_l = r_{l+1} - r_l \tag{1.2}$$

where $r_l$ is the coordinates of the Cα atom in residue $l$. The linking values $g_N(i,j)$ and $g_C(i,j)$ were calculated as:

$$g_{\mathrm{N}}(i,j) = \frac{1}{4\pi}\sum_{m=6}^{i-5}\sum_{n=i}^{j-1}\frac{R_m - R_n}{|R_m - R_n|^3}\cdot(dR_m \times dR_n) \tag{1.3}$$

$$g_{\mathrm{C}}(i,j) = \frac{1}{4\pi}\sum_{m=\mathrm{j}+6}^{\mathrm{N}-5}\sum_{n=i}^{j-1}\frac{R_m - R_n}{|R_m - R_n|^3}\cdot(dR_m \times dR_n) \tag{1.4}$$

where we excluded the first 5 residues on the N-terminal curve, last 5 residues on the C-terminal curve, and 4 residues before and after the native contact to eliminate the error induced by both the high flexibility and contiguity of those tails. A given loop is determined to be entangled if $|g_{\mathrm{N}}(i,j)| \geq 0.6$ or $|g_{\mathrm{C}}(i,j)| \geq 0.6$. If an NCLE is detected for a give native contact, we use the topoly[4] python package to identify the residues at which the termini cross the loop plane.

To control slipknots (which are rarely identified by this method but can occur due to the approximations made for a discrete structure with open curves) the chirality signs of the resulting crossings identified by Topoly[4] are used. If the sum of the chirality signs for a given terminus is 0 the NCLE with that terminus is disregarded. Also, if there is a crossing reported twice with different chirality's in the same NCLE, we disregard it as well as upon visual inspection they appear as artifacts caused by highly parallel crossing of the loop plane and uncertainty in the chirality.

##### 1.2. Filtering for high quality AlphaFold structures and NCLEs.

For AlphaFold structures with native NCLEs we have an additional set of criteria to reduce the inherent errors in the AF models from propagating to any downstream analysis.

1) The structure must have an overall <pLDDT> greater than or equal to 70. If not the whole structure is discarded.
2) The loop closing native contacts of an NCLE (i, j) must have a pLDDT >= 70.
3) The crossings must meet the following high quality order conditions:
    I. For a given termini that has an NCLE present start at the loop base and in order examine each crossing.
    II. If the first crossing has a pLDDT >= 70 it is kept and then you move onto the next.
    III. The first instance of a crossing where the pLDDT < 70 you discard this crossing and any other crossings after it. (if the first crossing after the loop base is low-quality, we disregard the whole NCLE with that temrini)

##### 1.3. Brief description of clustering NCLEs to obtain a representative NCLE

For each protein structure there are many degenerate NCLEs and therefore we seek to derive a set of unique representative NCLEs through a clustering algorithm provided in more detail in our previous publication[5]. As a high-level description this algorithm has three main steps:

1) Cluster together loops that share the same crossings identified by topoly and choose the smallest loop to be the representative loop of each subcluster.
2) Then merge subclusters based on meeting three criteria. - the crossings between the two sets of NCLEs is spatially close. (the details of this are complex and denoted in the cited paper) - loops overlap to any extent. - no crossing from either NCLE considered is within the loops of another in the cluster.
3) Finally, subclusters are merged based on whether the distance between their representative native contacts and crossings is less than a threshold defined by examining hand curated sets of NCLEs.

The result is for each protein a set of unique NCLE(s) is identified. For each unique NCLE a set of key entangled residues is defined as those residues within +/- 3 residues of the contacting residues $(i, j)$ or the crossing residues identified by topoly. The entangled region of each unique NCLE is then defined as the residues with an alpha carbon within 8 Å of the set of key entangled residues. Finally, we determine the overall entangled region of the protein to be the super set of the unique NCLE regions.

##### 1.4. Determining changes in proteolysis susceptibility across the proteome

To determine which sites across the proteome show a significant change in proteolysis susceptibility we examine the high throughput limited proteolysis mass spectrometry (LiP-MS) dataset previously published by the Fried group[7]. The proteome discover (PD) data was reanalyzed using FLiPPR[8] to determine unique probe protease (proteinase-K, PK) cut-sites that have significant changes in abundance between an untreated native sample and a sample refolded through treatment with 6M Guanidinium chloride and dilution jump into cytosol like medium. The FLiPPR protocol has the added benefits of an additional peptide merging step to combine peptides that report on the same PK cut site (probe protease) and filling of missing data in certain cases with gaussian imputations reflecting the mass spectrometry lower limit of detection. In brief the protocol is as follows:

1) Ions are initially separated into 4 classes:

1) Invalid peptide - not enough abundance in one or both cases (will be ignored in downstream analysis)

2) No missing abundances across all 6 samples

3) One abundance missing from either the Native or Refolded cases (missing value is dropped and statistical tests are preformed on the smaller sample set)

4) All-or-None: One sample is missing all 3 abundances while the other has all 3 (The 3 missing values are filled by Gaussian imputation of a distribution with mean and variance meant to mimic the lower limit of detection of the Mass spectrometry).

2) For class IV peptides the 3 missing values are filled by Gaussian imputation of a distribution with mean and variance meant to mimic the lower limit of detection of the Mass spectrometry ($\mu$ = 10,000 & $\sigma$ = 1,000).

3) If all the ions that inform on a given PK cut-site have abundance ratios that all agree in direction, then the overall ratio for the cut-site is calculated by taking the median of those ratios, and the P-values are combined with Fisher's method.

4) If there are two ions that inform on a given PK cut-site and they disagree, then a median is still calculated, but the P-value is set to 1. These cut-sites are also discounted from the pool of final peptides due to the uncertainty.

5) If there are three or more ions that inform on a given PK cut-site, then a majority rules heuristic is applied: the disagreeing ion is disregarded, and the (ratio, p-value)'s are only combined for the agreeing ions. If there is a tie, the P-value is set to 1. These cut-sites are still kept in the final pool of peptides.

Significant sites of change in proteolysis susceptibility for a given protein are then determined by the following criteria:

1. The unique PK cut-site has a statistically significant 2 fold increase or decrease in the abundance between the untreated sample (native) and a treated sample (refolded).
2. The statistical significance of the fold change is maintained after applying the BH false discovery rate correction to all the PK-cut-sites observed across the proteome (not just for the given protein).
3. The residue showed a change in proteolysis susceptibility at at-least one of the three timepoints after refolding (1min, 5min, 2hrs).

##### 1.5. Controlling for changes in protein abundance in statistical analysis of LiP-MS data

Controlling for effects of endogenous protein abundance differences in the LiP-MS data is done through both the sum of the peptide ion abundances (SPA) and the peptide coverage determined by PD. The SPA for protein $i$ is defined as:

$$SPA(i) \equiv \sum_{p} \langle a(p,i) \rangle \tag{5.1}$$

Where $\langle a(p,i) \rangle$ is the average peptide abundance reported by proteome discover for peptide $p$ in protein $i$ across the three untreated replicates. The SPA cumulative density function for each buffer (supplementary figure S1) is used to threshold groups of genes with a SPA greater than a given percentile. Every 10$^{th}$ percentile is considered to demonstrate robustness of our results to changes in protein abundance. For example, the 0% threshold considers all genes observed in the experiment for that buffer system regardless of abundance, while the 60% threshold considers those genes with a SPA greater than the 60$^{th}$ percentile of the data (i.e. the 40% most highly expressed). Finally only proteins that have a peptide coverage of 50% of greater of the canonical uniport sequence are considered in downstream analysis.

It is worth noting that protein length distribution after applying these cut offs does not significantly deviate from the natural full proteome length distribution.

##### 1.6. Association of non-refoldable proteins and native non-covalent lasso entanglements

We consider any protein observed in the limited proteolysis cyto-serum only dataset[7] that meet the abundance thresholds described in section 5 and have at least 1 peptide with a significant change in proteinase-K susceptibility as non-refoldable. All other proteins that also meet the abundance thresholds are considered refoldable. The ratio of the odds of a protein containing a native NCLE being non-refoldable to the odds of a protein containing a native NCLE being refoldable is then calculated from a 2x2 contingency table and the p-value for the significance of the association is calculated exactly using the Fisher Exact test. The odds ratio can also be mathematically defined by the following:

$$O.R. \equiv \frac{\left( P_{\mathrm{E,nR}} / P_{\mathrm{E,R}} \right)}{\left( P_{\mathrm{nE,nR}} / P_{\mathrm{nE,R}} \right)} \tag{6.1}$$

Where $P_{\mathrm{E,nR}}$ is the probability of a protein containing a native NCLE and not being refoldable, $P_{\mathrm{E,R}}$ is the probability of a protein containing a native NCLE and being refoldable, $P_{\mathrm{nE,nR}}$ is the probability of a protein not containing a native NCLE and not being refoldable, and $P_{\mathrm{nE,R}}$ is the probability of a protein not containing a native NCLE and being refoldable.

##### 1.7. Binomial logistic regression modeling log-odds of observing changes in proteolysis susceptibility across the proteome

We then fit a simple binomial logistic regression model using the statsmodel python package to estimate the log-odds of observing a significant change in proteolysis susceptibility as a function of the amino acid content and whether the residue was in the entangled region of the proteome or not.

$$\log\left(\frac{p}{1-p}\right) \approx \beta_0 + \beta_{\mathrm{region}} X_{\mathrm{region}} + \sum\nolimits_{AA} \beta_{AA} X_{AA} \tag{7.1}$$

Where $X_{\mathrm{region}}$ is a binary variable defining if the residue was in the entangled region of the protein or not, and $X_{AA}$ is a binary variable defining if the residue was of the type $AA$ where $AA$ is one of the 20 canonical amino acids. An odds ratio can then be defined as, $O.R. = e^{\beta_{\mathrm{region}}}$ where $\beta_{\mathrm{region}}$ if the resulting coefficient of the regression variable describing which region a given residue belonged in the binomial regression.

##### 1.8. Essential and non-essential gene definition

We define our set of essential genes as those identified in the knock-out experiments of Baba T and collaberators[9]. They identified 296 essential genes of which 188 are present in our set of high-quality crystal structures. We repeat the same binomial regression as described in the previous section for each group of genes and compare the resulting odd ratios to determine if indeed there is a dependance on gene essentiality.

##### 1.9. Propensity score matching of solvent accessible surface area in entangled regions of the proteome to non-entangled regions

To sample a set of residues from the non-entangled region of the proteome with similar solvent accessible surface area (SASA) to the entangled region we use the propensity score matching function in R on the set of per residue solvent accessible surface areas calculated on the set of 1294 high quality PDBs using the Shrake-Rupley method[10] in the MDtraj[11] python package with a probe radius of 1.4 Ǻ. We preform 1 to 1 matching, and the optimal caliper of 0.15 was chosen because it was the smallest caliper we could match all residues in the entangled region at least once and provided nearly identical PDFs for the entangled versus non-entangled region SASA. The binomial logistic regression described in the previous section was then refit to the propensity score matched dataset to determine if the resulting odds ratio significance changed.

##### 1.10. Scanning for DnaK binding motifs

Each protein in the set of 1294 we created three pools of sequences to search for potential DnaK binding sites (1) the whole FASTA sequence for the protein, (2) the loops sequences of each unique NCLE, and (3) the threads of each unique NCLE where the sequence for each thread was taken as the crossings that pierced the loop plane (as determined by topoly) +/- 10 residues. We then scanned each pool of sequences for matches to three different motif patterns derived from a combination of experiments[12], homology modeling[13], and an expanded Bukau-like motif that follow the pattern BHHHHHB, where B are basic residues and H are hydrophobic residues (Table S2.1).

The Bukau motifs were experimentally derived[12]. The Schymkowitz motif used a combination of experimentally derived motifs and structural parameters homology modeling to derive a more expansive position specific scoring matrix and we choose elements of that matrix with an enrichment score greater than 1 (i.e. greater than 10-fold the background)[13]. Finally, we choose to employ an Bukau-like motif which consists of all hydrophobic[14] residues flanked by basic residues to account for patterns observed across both Bukau and Schymkowitz motifs. As we are dealing with a limited dataset of proteins in the proteome and these motifs are often ensemble averages of sequences which DnaK was found to bind, we also examined the partial motifs were only the left flanking (LF) or right flanking (RF) basic residues and the hydrophobic core are included.

To compare the sets of essential and non-essential genes we calculate the number of motifs per 100 residues

$$D_i = \frac{N(i,p,m)}{L} * 100 \tag{10.1}$$

Where $i$ is the index of the gene, $p$ is the pool of sequences (i.e. whole sequence, just the loops, or just the threads), $m$ is the motif, $N(i,p,m)$ is then the number of hits for that motif in that pool for that gene, and $L$ is the length of the protein. The distributions of $D$ for each pool of genes can then be compared to determine if essential genes have more hits in entangled regions on average for DnaK binding sites than non-essential genes. Confidence intervals for these distributions were calculated by basic bootstrapping (n=10,000) and the test of a difference between the two distributions was done by permutation. Finally, we corrected the permutation p-values by the benjamini-hochberg procedure to control for false discovery rates.

##### 1.11. Non-covalent lasso entanglement complexity measures and feature selection to predict essential vs non-essential

For each protein with a native NCLE, we cluster the set of 'raw' native NCLEs into sets that represent the same topological NCLE and choose one with a minimal loop as the representative native NCLE for each set[5]. For each of these unique NCLEs we calculate 18 metrics designed to capture the complexity of a native NCLE, listed in the table below. For each set of proteins that are present in a given buffer condition (cyto-serum, +DnaK, and +GroEL) and meet a given SPA thresholded we estimate the boot-strapped confidence intervals (n=10,000) for the set of essential and non-essential genes and if there is a statistically significant difference in the distributions using the non-parametric Mann Whitney U test.

To determine which, if any, features best discriminate between the classes of essential and non-essential genes we use the Least Absolute Shrinkage and Selection Operator (LASSO) logistic regression model to find an optimal set of parameters. We scanned the inverse regularization strength $C$ from 0.00001 to 10 in 1,000 evenly placed steps. For each value of the penalty we fit the logistic regression model across 5 stratified k-folds of the standard scaled data in a 80:20 split. We then calculate the average balanced accuracy across the folds as a judge of predictive power. We also determine how many of the regression coefficients remained non-zero and had consistent sign across all folds as well to judge consistent robust results.

To determine if these balanced accuracies are significant beyond random chance we perform the LASSO regression method described above on randomly permuted versions of the dataset (n = 10,000) and determine what is the probability of observing an average balanced accuracy that is different from 0.5 by random chance. The optimal values of $C$ that provided the maximal balanced accuracy with the minimal number of non-zero robust features in the absence of chaperones was 1.5, in the presence of DnaK 2.5, and the presence of GroEL 2.5.

##### 1.12. Loop closing contact hydropathy and contact enrichment

To examine more details about which specific pairs of loop closing amino acids are enriched in essential or non-essential protein sets we calculate the odds ratio for the association of a given pair of amino acids $(a, b)$ being a loop closing contact and the protein being essential.

$$O.R.\left(s(B)\right) \equiv \frac{\left(N_{\mathrm{lc}}^{\mathrm{nE}}(a,b)\right)/\left(N_{\mathrm{lc}}^{\mathrm{nE}} - N_{\mathrm{lc}}^{\mathrm{nE}}(a,b)\right)}{\left(N_{\mathrm{lc}}^{\mathrm{E}}(a,b)\right)/\left(N_{\mathrm{lc}}^{\mathrm{E}} - N_{\mathrm{lc}}^{\mathrm{E}}(a,b)\right)} \quad (12.1)$$

Where $s(B)$ is the set of proteins that have a SPA greater than the $50^{\mathrm{th}}$ SPA percentile and have more than 50% of their primary sequence observed in the LiP-MS experiments in buffer condition $B$. $N_{\mathrm{lc}}^{\mathrm{nE}}$ and $N_{\mathrm{lc}}^{\mathrm{E}}$ are the number of loop-closing contacts in the set of non-essential and essential proteins respectfully. $N_{\mathrm{lc}}^{\mathrm{nE}}(a,b)$ and $N_{\mathrm{lc}}^{\mathrm{E}}(a,b)$ is the number of loop-closing contacts composed of residue types $(a, b)$ in the set of non-essential and essential proteins respectfully.

We determine which pairs of amino acids $a$ & $b$ show a statistically significant enrichment in either the non-essential or essential protein classes by permutation of the loop-closing contacts (n = 100,000). After false discovery rate correction using the benjamini-hochberg procedure we consider the significant pairs to have an adjusted p-value less than 0.05 in all three buffer conditions (cyto-serum only, +DnaK, and +GroEL) to ensure we are only considering the enrichment of the most robust pairs of loop closing amino acids observed across all experiments.

##### 1.13. Selecting candidates for coarse grained molecular dynamics simulations

Three sets of proteins are selected for simulation and screening of misfolding mechanisms. The first dataset (I) is the set of proteins that are predicted to show minimal too no conformational changes unpon refolding. They have no significant observed changes in abundance of any half-tryptic or full tryptic peptides in the chaperone free 5min and 2hr post refolding LiP-MS datasets.

Candidate sets (II) and (III) are chosen from pools of proteins with a high likelihood of misfolding involving a native NCLE. These pools are generated through Monte Carlo style simulated annealing trajectories launched from random distribution of initial candidates selected by the following criteria:

1) have atleast 50% of their canonical sequence observed in the chaperone free LiP-MS experiments
2) have a sum of their peptide abundance (SPA) greater than the 50$^{th}$ SPA percentile of all proteins in the chaperone free conditions.
3) be non-refoldable in the absence of chaperones (i.e. have at least 1 significant PK site)

These initial candidates are randomly distributed amongst 4 groups of equal size. For each group of proteins $i$ there is an objective function defined as:

$$E_i = -C_1\, log(OR_i) + C_2\, D_i \tag{13.1}$$

Where $OR_i$ is the ratio of the odds of misfolding in the entangled region relative to the odds of misfolding in the non-entangled region of the protein group $i$. Where $D_i$ is the Kolmogorov–Smirnov test statistic for the protein size distribution and the reference distribution taken as the superset of all four groups. One MC step is taken by randomly swapping 5 proteins between each pair of neighboring group indexes. $C_1 = 1$ and $C_2 = 2.5$ in this work and was optimized to ensure penalties for deviations from the reference protein size distribution are balanced by the opportunity to increase odds ratio.

The Metropolis criteria[15] is applied to determine if any given swap is accepted or rejected. Where the acceptance ratio for a swap between groups $j$ and $k$ at MC step $x$ is defined as:

$$P \equiv \frac{P_{\text{new}}(j,k,x)}{P_{\text{old}}(j,k,x-1)} \propto e^{-\beta\Delta E} = e^{-\beta(E_{\text{new}}-E_{\text{old}})} = e^{-\beta\left(E_j^x + E_k^x - E_j^{x-1} - E_k^{x-1}\right)} \tag{13.2}$$

The swap is accepted if either $P \geq 1$ or $1 > P > u$, where $u$ is a random number sampled from a uniform distribution bounded by [0,1]. β is an inverse temperature scaling factor that starts very low at 0.05 to allow ample room for the simulation to explore the objective function landscape and is scaled linearly to 1000 after every 750 MC steps and remains constant at 1000 thereafter. Plots of the energy derivative as a function of temperature show a clear phase transition observed over this window.

In total 10 independent trajectories are run for the set of proteins with native NCLEs that are likely to be non-refoldable. A parallel set of 10 independent simulations was run with a randomized proteome sample where the significant PK cut-sites are distributed randomly across the proteome. This later set acts as a randomized control we use to measure the inherent shuffling of noise present in these simulations.

Candidates set we predict are highly likely to misfold involving their native NCLE, (II) and (III), were selected through rank ordering the proteins by their fractional presence in the group with the largest OR at the end of each trajectory, $f_M \geq 0.7$. The largest OR group is defined as the group with the highest mean OR over the last 10 frames of the trajectory. Candidates for set II were chosen to match the size distribution of set I as close as possible. Candidates for set III were chosen at random to reflect the underlying distribution.

##### 1.14. Coarse graining simulation candidates

In this study, we employed a Gō-based force field for all proteins analyzed, as described in prior work[16–20]. Briefly, this coarse-grained (CG) model represents each residue as a single interaction site centered at the Cα atom position. The potential energy function is given by:

$$\begin{aligned} E_{\text{tot}} &= E_{\text{bond}} + E_{\text{angle}} + E_{\text{dihedral}} + E_{\text{elec}} + E_{\text{vdW}} \\ &= \sum_i K_{\text{b}}(b_i - b_0)^2 + \sum_i -\frac{1}{\gamma}\ln\left\{e^{-\gamma[K_\alpha(\theta_i-\theta_\alpha)^2+\varepsilon_\alpha]} + e^{-\gamma K_\beta(\theta_i-\theta_\beta)^2}\right\} \\ &\quad + \sum_i \sum_{j=1}^{4} K_{\text{D}_j}\left[1+\cos\left(j\varphi_i - \delta_j\right)\right] + \sum_{i,j} \frac{q_i q_j}{4\pi\varepsilon_0\varepsilon_\gamma r_{ij}} \cdot e^{-\frac{r_{ij}}{l_{\text{D}}}} \\ &\quad + \sum_{i,j} \varepsilon_{ij}\left[13\left(\frac{R_{ij}}{r_{ij}}\right)^{12} - 18\left(\frac{R_{ij}}{r_{ij}}\right)^{10} + 4\left(\frac{R_{ij}}{r_{ij}}\right)^{6}\right] \end{aligned} \tag{14.1}$$

Here, $K_b$ is the bond force constant, set to 50 kcal/mol/Å22; $b_i$ represents the $i$-th pseudo-bond length between two adjacent interaction sites, with $b_0$ as the equilibrium pseudo-bond length, fixed at 3.81 Å, the average distance between adjacent Cα atoms in the protein sequence. Constants $\gamma, K_\alpha, \theta_\alpha, \varepsilon_\alpha, K_\beta$ , and $\theta_\beta$ define the double-well angle potential[21], which accounts for bond angles associated with both α-helix and β-sheet conformations. $\theta_i$ is the $i$-th bond angle, while $K_{\text{D}_j}$ and $\delta_j$ represent the dihedral force constants and phases for periodicity $j$, respectively, with $\varphi_i$ as the $i$-th pseudo-dihedral angle.

The charge $q_i$ corresponds to the net charge of the $i$-th interaction site, matching the net charge of the corresponding amino acid residue. $\varepsilon_0$ and $\varepsilon_\gamma$ denote the dielectric constants of vacuum and water, respectively. The Debye length, $l_D$, is set to 10 Å. The Lennard-Jones (LJ)[22] potential parameters, $\varepsilon_{ij}$ and $R_{ij}$, account for desolvation barriers between interaction sites $i$ and $j$, with $r_{ij}$ as their distance. All 1-4 nonbonding interactions are included without scaling any parameters. Nonbonding interactions are smoothly tapered to zero between 18 Å and 20 Å. Bond length constraints are enforced throughout all simulations.

For sites forming native contacts, $\varepsilon_{ij} = \varepsilon_{ij}^{HB} + \eta_{\text{scale}}\varepsilon_{ij}^{SC-SC} + \varepsilon_{ij}^{BB-SC}$. The hydrogen bond (HB) interaction depth $\varepsilon_{ij}^{HB}$ is 0.75 kcal/mol for single HB contacts and 1.5 kcal/mol for multiple HB contacts. The sidechain-sidechain (SC-SC) interaction depth $\varepsilon_{ij}^{SC-SC}$ is derived from the Betancourt−Thirumalai statistical potential[23] and scaled by $\eta_{\text{scale}}$ to achieve realistic native-state stability for each protein. The backbone-sidechain (BB-SC) interaction depth $\varepsilon_{ij}^{BB-SC}$ is set at 0.37 kcal/mol. Native contacts are defined for residue pairs separated by at least two residues in the native structure. STRIDE[6] identifies native HB contacts, while SC-SC and BB-SC contacts are identified as residue pairs with any heavy atom within 4.5 Å in the crystal structure. The value of $R_{ij}$ for native contacts is set as the native distance $d_{ij}$ between sites $i$ and $j$.

For non-native contacts, the interactions are primarily repulsive, with $\varepsilon_{ij} = \sqrt{\varepsilon_i \varepsilon_j}$ and $R_{ij} = R_i + R_j$ , where $\varepsilon_i$ is set to 0.000132 kcal/mol, and $R_i$ is the non-native collision diameter $\sigma_i$ of site $i$, multiplied by $2^{1/6}$ and divided by 2. These collision diameters are determined based on the protein's native structure using the Karanicolas−Brooks method[22]. Simulations were performed in OpenMM[24].

##### 1.15. Optimizing $n_{\text{scale}}$ for candidates

To parameterize proteins more complex than those in our training set, we employed a stepwise optimization strategy. We predefined five levels of $\eta_{\text{scale}}$ for each structural class, as detailed in Supplementary Table S2.6. For more details on how these values were derived see our previous publications[25,26], but in brief here is how the levels are defined. First Level: At this level, $\eta_{\text{scale}}$ was set to the mean value for the structural class, $\langle \eta_{\text{scale}}^* \rangle_{class}$, as well as the overall mean value of the 18 training-set proteins for the domain interface. Second Level: Here, $\eta_{\text{scale}}$ is adjusted to $\langle \eta_{\text{scale}}^* \rangle_{class}$ plus an increase of $\langle \frac{\Delta G_{\text{UN}}^{\text{exp}} - \Delta G_{\text{UN}}^{class}}{\Delta G_{\text{UN}}^{class}} \rangle_{-} \cdot 100 = 23\%$, where $\Delta G_{\text{UN}}^{class} = m \cdot \langle \eta_{\text{scale}} \rangle_{class} + b$. The notation $\langle \ldots \rangle_{-}$ means the average is only calculated for the proteins that are destabilized using $\langle n_{\text{scale}}^* \rangle_{class}$ within the training set. Third Level: $\eta_{scale}$ was further increased to $\langle \eta_{\text{scale}}^* \rangle_{class}$ plus an increase of $max \left[ \frac{\Delta G_{\text{UN}}^{\text{exp}} - \Delta G_{\text{UN}}^{class}}{\Delta G_{\text{UN}}^{class}} \right]_{-} \cdot 100 = 46\%$, where the maximum is taken only for proteins destabilized using $\langle \eta_{\text{scale}}^* \rangle_{class}$ in the training set. Fourth Level: $\eta_{\text{scale}}$ was increased by $\frac{min(\Delta G_{\text{UN}}^{\text{exp}} - \Delta G_{\text{UN}}^{class})}{\langle \Delta G_{\text{UN}}^{\text{class}} \rangle} \times 100\% = 70\%$ representing the largest destabilization relative to the overall average $\Delta G_{\text{UN}}^{class}$. Fifth Level: Finally, $\eta_{\text{scale}}$ was set to the overall maximum value across all levels.

We parameterized an arbitrary protein using the following strategy: First, the protein was coarse-grained. Native van der Waals (vdW) interactions were grouped based on domains defined by CATH[27], and $\eta_{\text{scale}}$ was initially assigned using the first-level value according to the domain's structural class (or interface).

Next, we performed 10 parallel 0.5-μs MD simulations at 310 K for the coarse-grained model, monitoring the fraction of native contacts ($Q$) of each domain and interface. A domain or interface was considered stabilized if all 10 trajectories showed $Q$ values greater than the threshold $\langle Q_{\text{kin}} \rangle = 0.6688$ for at least 98% of the simulation time. For destabilized domains or interfaces, $\eta_{\text{scale}}$ was increased to the next level, while stabilized regions retained their $\eta_{\text{scale}}$ values, and a new CG model was generated. This process was repeated until all domains and interfaces were stabilized.

If a domain or interface could not be stabilized even at the highest $\eta_{\text{scale}}$ level, the median $\eta_{\text{scale}}$ value for the corresponding structural class was used to generate the final CG model, regardless of stability.

##### 1.16. Temperature quenching simulations of 28 candidate proteins

For each of the candidates in supplementary table S2.6 we run 50 independent 1 us unfolding simulations at 800 K for their coarse-grained models using Langevin dynamics in OpenMM[24] with a 0.05 $ps^{-1}$ friction coefficient. Frames are saved every 5000 steps with a timestep of a 0.015 ps.

A trajectory was considered unfolded when the mode of the fraction of native contacts ($Q_{\text{mode}}$) over the previous 100ns of the simulation was less than 0.15 for 100 frames. Once the condition was met the trajectory was halted and the final frame was saved as the starting unfolded structure. For each candidate the 50 unfolded structures were quenched instantaneously to 310 K and then another 2 us of Langevin dynamics were run.

##### 1.17. Clustering changes in non-covalent lasso entanglement in temperature quenching trajectories

For each frame $f$ of a trajectory $t$ we then cluster the changes in NCLE observed across the set of all loops $\Gamma(f) \equiv \{\gamma_1, \dots, \gamma_k\}$ where $\gamma_k$ is the $k^{th}$ loop identified to have a change in its rounded Gauss linking value ($G_{\text{n}}$ and $G_{\text{c}}$) between the frame $f$ and the reference structure. Please note we round based on a threshold of 0.6 instead of the typical 0.5 to help mitigate false positives in the identification of a loss or gain of NCLE. First, we remove any changes observed in either the N or C terminus for any given loop in the frame that have an absolute value change in the Gauss linking value between the frame and the reference structure less than 0.35. This is to control for phantom changes in NCLE that are introduced by the required threshold to identify a loop and thread combination as entangled (defined in the "Determining unique NCLEs across the set of 1294 representative protein structures" section). We also do not consider pure switches in chirality in this work as they are relegated to ~6% of the changes in NCLE observed, do not change the overall conclusions, and are highly suspect as to whether they are artifacts of our coarse-grained model. We then group the changes based on the following clustering vector:

$$\boldsymbol{v}(f) \equiv \left[G_{\text{n}}, \Delta_{\text{n}}, \left|N_{\text{crossings/n}}\right|, G_{\text{c}}, \Delta_{\text{c}}, \left|N_{\text{crossings/c}}\right|\right] \quad (17.1)$$

Where $G_{\text{n}}$ and $G_{\text{c}}$ are the rounded Gauss linking numbers for the N and C terminus respectively. $\Delta_{\text{n}}$ and $\Delta_{\text{c}}$ are the type of change observed in either the N or the C terminus which can fall into 5 categories: Loss of a native NCLE, loss of a native NCLE with a switch in chirality, gain of a native NCLE, gain of a native NCLE with a switch in chirality, or a pure switch in chirality. These categories are defined more robustly in the SI of the following references[25,28]. $\left|N_{\text{crossings/n}}\right|$ and $\left|N_{\text{crossings/c}}\right|$ are the number of crossings observed in the new non-native NCLE if the change is of a gain nature (gain of a native NCLE, gain of a native NCLE with a switch in chirality), or the number of crossings observed in the reference native state structure if the change is of a loss nature (loss of a native NCLE, loss of a native NCLE with a switch in chirality). All elements of $v(f)$ are integers and thus as the initial clustering step is concerned, we require all members of the cluster $v_c(f)$ to have a Euclidian distance to any other member of 0.

Next within each cluster $v_c(f)$ we then cluster based on the Euclidean distance between the median crossing residues of each $\gamma_k$. Again, if the change was off a gain nature the crossings in the frame $f$ are used and if the change is of a loss nature we use the crossings in the reference structure for the given loop and thread. We use the DBSCAN[29] clustering method with a maximum neighborhood distance of 10 and a minimum number of samples to be considered a neighborhood of 1 to define subclusters in $v_c(f)$.

Finally, within each of these crossing subclusters clusters we cluster the loops based on their loop closing contact residue indexs $(i, j)$. We use a distance formula that aims to cluster based on the overlap between any two loops $\gamma_k$ and $\gamma_h$.

$$d_{\text{loop}} = \frac{\max(\gamma_k \cup \gamma_h) - \min(\gamma_k \cup \gamma_h)}{|\gamma_k| + |\gamma_h|} \tag{17.2}$$

The distance equals 1.0 when two loops are adjacent, becomes less than 1.0 when they overlap, and greater than 1.0 when they are distant. We again employ DBSCAN using a maximum neighborhood distance of 1 and a minimum number of samples to be considered a neighborhood of 1 resulting in the final set of clustered unique changes in NCLE.

##### 1.18. Calculating probability of misfolding mechanism type

To calculate the probability of observing either a loss of a native NCLE or a gain of a non-native NCLE we first filter the trajectories remove those that are natively folded in the last 200 ns of trajectory $t$ of a protein $i$ in candidate set (III) (defined in the "Selecting candidates for coarse grained molecular dynamics simulations" section). Trajectories are considered natively folded and discarded if $Q^{\text{mode}}_{(200\text{ ns})} \geq Q_{\text{threshold}}$ and $G^{\text{mode}}_{(200\text{ ns})} \leq G_{\text{threshold}}$, where $Q_{\text{threshold}}$ and $G_{\text{threshold}}$ thresholds are set by the reference simulations (Equations (8) and (9), supplementary tables 2.7 - 2.8). For those trajectories that pass the filtering criteria we calculate the average lifetime of the misfolded state in each trajectories as the average fraction of consecutive frames where a change of NCLE status was identified multiplied by 0.075 ns as the timestep. We then separate the trajectories into long lived and short-lived misfolded states based on the median lifetime of ~100 ns.

To calculate the probability of a given structure exhibiting a misfolding mechanism $M$ (only a loss of a native NCLE, only a gain of a non-native NCLE, or both) within each trajectory $t$ we use the following:

$$P(M) = \frac{1}{N_{t,i}} \sum_{t,i} \frac{N_M(t,i)}{N_{\text{only-loss}}(t,i) + N_{\text{only-gain}}(t,i) + N_{\text{both}}(t,i)} \tag{18.1}$$

Where $N_M(t,i)$ is the number of structures observed in the quenching trajectory $t$ of protein $i$ with the particular misfolding type $M$ present and $N_{t,i}$ is the total number of trajectories across all protein in a given dataset considered. We then report the ensemble average of $P(M,t,i)$ across all proteins and quenching trajectories that met the criteria above.

Considering only the structures that contain both a loss of a native NCLE and the gain of a non-native NCLE we calculate the conditional probability that a loss is coupled (or paired) with a gain in a given trajectory $t$ by examining whether the key topological features (i.e. the loop closing native contacts +/- 3 residues and the crossing residues +/- 3 residues) of the NCLEs are shared. The conditional probability of a loss of a native NCLE being paired with at least 1 gain of a non-native NCLE in the last 200 ns of trajectory $t$ of protein $i$ is then given the structure had both present is then defined as:

$$P = \frac{1}{N_{t,i}} \sum_{t,i} \frac{N_{\mathrm{p}}(t,i)}{N_{\mathrm{loss}}(t,i)} \tag{18.2}$$

Where $N_{\mathrm{p}}(t,i)$ is the number of losses of NCLE that had primary structure overlap of key NCLE topological features with at least 1 gain of NCLE in the same structure, $N_{\mathrm{loss}}(t,i)$ is the total number of losses observed across all structures containing both a types of misfolding, and $N_{t,i}$ is the total number of trajectories across all protein in a given dataset considered.

### 2. Supplemental data tables

#### Supplementary Table 2.1 | DnaK binding motif definitions

| | Position along primary structure | | | | | | |
|---|---|---|---|---|---|---|---|
| **Motif** | **0** | **1** | **2** | **3** | **4** | **5** | **6** |
| Bukau* | R, K | I, L, V, F, Y | I, L, V, F, Y | I, L, V, F, Y | I, L, V, F, Y | R, K | - |
| Bukau(LF)* | R, K | I, L, V, F, Y | I, L, V, F, Y | I, L, V, F, Y | I, L, V, F, Y | - | - |
| Bukau(RF)* | - | I, L, V, F, Y | I, L, V, F, Y | I, L, V, F, Y | I, L, V, F, Y | R, K | - |
| Bukau-like** | R, K, H | I, L, V, F, Y, A, M, W | I, L, V, F, Y, A, M, W | I, L, V, F, Y, A, M, W | I, L, V, F, Y, A, M, W | R, K, H | - |
| Bukau-like(LF)** | R, K, H | I, L, V, F, Y, A, M, W | I, L, V, F, Y, A, M, W | I, L, V, F, Y, A, M, W | I, L, V, F, Y, A, M, W | - | - |
| Bukau-like(RF)** | - | I, L, V, F, Y, A, M, W | I, L, V, F, Y, A, M, W | I, L, V, F, Y, A, M, W | I, L, V, F, Y, A, M, W | R, K, H | - |
| Schymkowitz*** | K, E, Q, W, Y | F, I, K, L, R, V, Y | F, L, R, V, W, Y | I, L, M, T, V | F, L, M, P, R, V, Y | F, I, L, W, Y | F, N, P, R, Y |

* motif reported in reference 12
** motif reported in reference 12 using an expanded set of hydrophobic and basic residues types
*** motif reported in reference 13
(LF) keeping the left flanking basic residues but excluding the right flanking
(RF) keeping the right flanking basic residues but excluding the left flanking

#### Supplementary Table 2.2 | DnaK binding motif hit rate per 100 residue statistics

| Motif | Pool type | Essential proteins | | | Non-essential proteins | | | ratio | \|difference\| | p-value |
|---|---|---|---|---|---|---|---|---|---|---|
| | | <D> | 95% ci lower | 95% ci upper | <D> | 95% C.I. lower | 95% C.I. upper | | | |
| Bukau* | All | 0.0165 | 0.0000 | 0.0631 | 0.0022 | 0.0000 | 0.0131 | 7.5631 | 0.0143 | 0.7921 |
| | Thread | 0.0000 | 0.0000 | 0.0000 | 0.0022 | 0.0000 | 0.0109 | 0.0000 | 0.0022 | 1.0000 |
| | Loop | 0.0058 | 0.0000 | 0.0290 | 0.0000 | 0.0000 | 0.0000 | inf | 0.0058 | 0.8705 |
| Bukau(RF)* | All | 0.0312 | 0.0119 | 0.0758 | 0.0283 | 0.0139 | 0.0532 | 1.1025 | 0.0029 | 1.0000 |
| | Thread | 0.0104 | 0.0000 | 0.0358 | 0.0109 | 0.0041 | 0.0249 | 0.9551 | 0.0005 | 1.0000 |
| | Loop | 0.0131 | 0.0014 | 0.0422 | 0.0152 | 0.0066 | 0.0302 | 0.8629 | 0.0021 | 1.0000 |
| Bukau(LF)* | All | 0.0846 | 0.0445 | 0.1500 | 0.0605 | 0.0412 | 0.0877 | 1.3976 | 0.0241 | 0.8705 |
| | Thread | 0.0197 | 0.0060 | 0.0498 | 0.0188 | 0.0099 | 0.0316 | 1.0472 | 0.0009 | 1.0000 |
| | Loop | 0.0396 | 0.0177 | 0.0835 | 0.0262 | 0.0151 | 0.0430 | 1.5122 | 0.0134 | 0.8705 |
| Schymkowitz*** | All | 0.0144 | 0.0021 | 0.0439 | 0.0058 | 0.0022 | 0.0130 | 2.4693 | 0.0085 | 0.8705 |
| | Thread | 0.0050 | 0.0000 | 0.0269 | 0.0013 | 0.0000 | 0.0064 | 3.8990 | 0.0037 | 0.8705 |
| | Loop | 0.0050 | 0.0000 | 0.0299 | 0.0033 | 0.0010 | 0.0093 | 1.5004 | 0.0017 | 1.0000 |
| Bukau-like** | All | 0.0632 | 0.0316 | 0.1211 | 0.0399 | 0.0262 | 0.0583 | 1.5848 | 0.0233 | 0.8705 |
| | Thread | 0.0009 | 0.0000 | 0.0044 | 0.0139 | 0.0065 | 0.0263 | 0.0634 | 0.0130 | 0.7921 |
| | Loop | 0.0215 | 0.0093 | 0.0457 | 0.0236 | 0.0136 | 0.0394 | 0.9091 | 0.0021 | 1.0000 |
| Bukau-like(RF)** | All | 0.2844 | 0.2228 | 0.3654 | 0.3241 | 0.2783 | 0.3729 | 0.8776 | 0.0397 | 0.8705 |
| | Thread | 0.0374 | 0.0172 | 0.0801 | 0.0899 | 0.0655 | 0.1221 | 0.4160 | 0.0525 | 0.6728 |
| | Loop | 0.1329 | 0.0929 | 0.1855 | 0.1430 | 0.1139 | 0.1787 | 0.9292 | 0.0101 | 1.0000 |
| Bukau-like(LF)** | All | 0.3605 | 0.2932 | 0.4448 | 0.3981 | 0.3497 | 0.4511 | 0.9055 | 0.0376 | 0.8705 |
| | Thread | 0.0732 | 0.0438 | 0.1172 | 0.0933 | 0.0714 | 0.1216 | 0.7852 | 0.0200 | 0.8705 |
| | Loop | 0.1513 | 0.1058 | 0.2074 | 0.1810 | 0.1468 | 0.2246 | 0.8358 | 0.0297 | 0.8705 |

* motif reported in reference 12
** motif reported in reference 12 using an expanded set of hydrophobic and basic residues types
*** motif reported in reference 13
(LF) keeping the left flanking basic residues but excluding the right flanking
(RF) keeping the right flanking basic residues but excluding the left flanking

#### Supplementary Table 2.3 | Definitions of 18 topological complexity descriptors for native non-covalent lasso entanglements

| Feature | Feature ID | Description |
|---|---|---|
| $g_n$ | 0 | $g_N(i,j) = \frac{1}{4\pi}\sum_{m=6}^{i-5}\sum_{n=i}^{j-1}\frac{R_m - R_n}{\|R_m - R_n\|^3}\cdot(dR_m \times dR_n)$ |
| $N_{\text{Threads}}$ | 1 | Number of N-terminal thread crossings of the loop plane (determined) by the topoly python package |
| $g_c$ | 2 | $g_C(i,j) = \frac{1}{4\pi}\sum_{m=j+6}^{N-5}\sum_{n=i}^{j-1}\frac{R_m - R_n}{\|R_m - R_n\|^3}\cdot(dR_m \times dR_n)$ |
| $C_{\text{Threads}}$ | 3 | Number of C-terminal thread crossings of the loop plane (determined) by the topoly python package |
| $N_{\text{Loop}}$ | 4 | The size of the representative NCLE loop enclosed by the native contact $(i,j)$ |
| $N_{\text{Zipper}}$ | 5 | The number of loop closing native contacts encompassing the same NCLE that form the cluster from which the representative NCLE is chosen. (contacts defined by any two residues that have heavy atoms within 4.5Å and are at least 3 residues apart along the primary structure) |
| $f_{\text{BB,Loop}}$ | 6 | The percent of the protein primary sequence that the loop encompasses |
| $N_{\text{Loop-cont}}$ | 7 | The sum of the number of contacts each residue in the loop makes with the rest of the protein. (contacts defined by any two residues that have alpha carbons within 8Å and are at least 3 residues apart along the primary structure) |
| $N_{\text{Cross-cont}}$ | 8 | The sum of the number of contacts each crossing residue(s) and buffer(s) (buffer defined as +/- 3 residues along the primary sequence) makes with the rest of the protein (contacts defined by any two residues that have alpha carbons within 8Å and are at least 3 residues apart along the primary structure) |
| $f_{\text{entR}}$ | 9 | The percentage of the protein defined as an entangled region where an entangled region defined as follows. For each protein's unique NCLE(s) identified we define a set of key loop closing residues as those residues within +/- 3 residues of the contacting residues $(i,j)$. Similarly, for the threading residues of each unique NCLE we identify a set of key residues that are within +/- 3 residues of the thread residues that cross the loop plan as determined by topoly. We then define the entangled region of each unique NCLE as the residues with an alpha carbon within 8Å of the set of key loop closing and threading residues. |
| $\text{MIN}[d_{\text{P,N}}]$ | 10 | Depth of the N-terminal crossings from the N-terminus (in residues), normalized by the length of the protein $\frac{n}{L_P}$ |
| $\text{MIN}[d_{\text{T,N}}]$ | 11 | Depth of the N-terminal crossings from the N-terminus (in residues), normalized by the length of the N-terminal thread $\frac{n}{i}$ |
| $\text{MIN}[d_{\text{S,N}}]$ | 12 | Depth of the N-terminal crossings from the N-terminus (in residues) $n$ |
| $\text{MIN}[d_{\text{P,C}}]$ | 13 | Depth of the C-terminal crossings from the C-terminus (in residues), normalized by the length of the protein $\frac{L_P - c}{L_P}$ |
| $\text{MIN}[d_{\text{T,C}}]$ | 14 | Depth of the C-terminal crossings from the C-terminus (in residues), normalized by the length of the thread $\frac{L_P - c}{L_P - j}$ |
| $\text{MIN}[d_{\text{S,C}}]$ | 15 | Depth of the C-terminal crossings from the C-terminus (in residues) $L_P - c$ |
| $ACO$ | 16 | Absolute contact order defined as $\frac{1}{N_{\text{Zipper}}}\sum_{c \in C_{GE}}(j-i)$ Where, $(i,j)$ are the loop closing native contact residue ids. $C$ is the set of loop closing native contacts encompassing the same NCLE that form the cluster from which the representative NCLE is chosen |
| $RCO$ | 17 | Relative contact order defined as $\frac{ACO}{L_P}$ |

* Code for computing these metrics is available on this manuscripts associated git-hub repository

#### Supplementary Table 2.4 | Associations between native non-covalent lasso entanglements, changes in proteolysis susceptibility, and protein essentiality is consistent between experimental and AlphaFold datasets

| | Description | Buffer | Essentiality | Crystal structures | | High quality AlphaFold structures | |
|---|---|---|---|---|---|---|---|
| | | | | Odds Ratio | p-value | Odds Ratio | p-value |
| (i) | Association of native NCLEs and non-refoldability | cyto-serum | - | 3.06 (1.55, 6.02) | $2.9x10^{-6}$ | 2.42 (1.47, 3.92) | $4.86x10^{-4}$ |
| (ii) | entangled regions of proteins are more likely to misfold than their non-entangled regions | cyto-serum | - | 1.44 (1.31, 1.58) | $3.10x10^{-13}$ | 1.34 (1.24, 1.45) | $4.66x10^{-12}$ |
| | entangled regions of proteins are more likely to misfold than their non-entangled regions even after controlling for the potential confounding factors of expression level and residue burial | cyto-serum | - | 1.38 (1.24, 1.54) | $6.79x10^{-9}$ | 1.33 (1.21, 1.46) | $5.47x10^{-9}$ |
| (iii) | that DnaK and GroEL do not, on average, correct misfolding of the entangled regions relative to the non-entangled regions | +DnaK | - | 1.27 (1.02, 1.57) | 0.026 | 1.43 (1.19, 1.72) | $1.33x10^{-4}$ |
| | | +GroEL | - | 1.22 (1.035, 1.45) | 0.018 | 1.33 (1.15, 1.54) | $1.348x10^{-4}$ |
| (iv) | that essential and non-essential show equal amounts of native NCLE misfolding | cyto-serum | Essential | 1.46 (1.22, 1.74) | $3.25x10^{-5}$ | 1.47 (1.26, 1.70) | $4.75x10^{-7}$ |
| | | | Non-essential | 1.43 (1.27, 1.61) | $2.41x10^{-9}$ | 1.30 (1.18, 1.43) | $1.17x10^{-7}$ |
| (v) | that the chaperones rescue the misfolding of essential proteins but not non-essential proteins | +DnaK | Essential | 1.10 (0.74, 1.64) | 0.624 | 1.15 (0.82, 1.63) | 0.404 |
| | | | Non-essential | 1.35 (1.05, 1.74) | 0.021 | 1.56 (1.26, 1.94) | $6.49x10^{-5}$ |
| | | +GroEL | Essential | 1.04 (0.75, 1.43) | 0.804 | 1.06 (0.81, 1.39) | 0.6599 |
| | | | Non-essential | 1.31 (1.06, 1.59) | 0.009 | 1.47 (1.23, 1.75) | $2.35x10^{-5}$ |
| (vi) | The association of protein essentiality with being a known GroEL client | cyto-serum | - | 0.95 (0.52, 1.61) | 0.89 | 1.34 (0.82, 2.16) | 0.26 |
| | The association of protein essentiality with being a known DnaK client | cyto-serum | - | 1.62 (0.94, 2.90) | 0.10 | 1.72 (1.07, 2.87) | 0.03 |

#### Supplementary Table 2.5 | Loop forming contact enrichment in non-essential proteins is consistent between experimental and AlphaFold datasets

| | | Crystal structures | | High quality AlphaFold structures | |
|---|---|---|---|---|---|
| **Buffer** | **Contact** | **Odds Ratio** | **p-value** | **Odds Ratio** | **p-value** |
| cyto-serum | S-Y | 4.97 | 0.0002 | 2.39 | 0.0014 |
| | H-C | inf | < 0.0001 | inf | < 0.0001 |
| | Y-F | 3.15 | < 0.0001 | 2.28 | < 0.0001 |
| | F-F | 7.13 | < 0.0001 | 7.44 | < 0.0001 |
| +DnaK | S-Y | 8.57 | 0.0004 | 3.45 | 0.0005 |
| | H-C | inf | < 0.0001 | inf | < 0.0001 |
| | Y-F | 3.52 | 0.0005 | 3.21 | < 0.0001 |
| | F-F | 3.51 | 0.0044 | 5.30 | < 0.0001 |
| +GroEL | S-Y | 5.90 | 0.0003 | 2.71 | 0.0023 |
| | H-C | inf | < 0.0001 | inf | < 0.0001 |
| | Y-F | 4.02 | < 0.0001 | 2.50 | < 0.0001 |
| | F-F | 5.06 | 0.0002 | 6.04 | < 0.0001 |

#### Supplementary Table 2.6 | Optimized η-scale values of coarse-grained models of simulation candidates

| Gene | PDB | Chain | Length | ID | resid | Domain Type(s) | η-scale |
|---|---|---|---|---|---|---|---|
| P0A7N9 | 6XZ7 | b | 55 | 0 | 1:55 | beta | 2.5044 |
| P0AG51 | 6XZ7 | Z | 59 | 0 | 1:59 | alpha-beta | 1.9644 |
| P0ADZ0 | 6QUL | U | 100 | 0 | 1:100 | alpha-beta | 1.9644 |
| P0AA25 | 6LUR | D | 109 | 0 | 1:109 | beta | 1.4732 |
| P61175 | 6XZ7 | S | 110 | 0 | 1:110 | alpha-beta | 1.6871 |
| P77214 | 1ZEQ | X | 110 | 0 | 1:110 | beta | 1.812 |
| P0ADY3 | 6XZ7 | K | 123 | 0 | 1:123 | beta | 1.4732 |
| P0A790 | 4CRZ | A | 126 | 0 | 1:126 | beta | 1.812 |
| P0A6T9 | 3A7L | A | 129 | 0 | 1:129 | beta | 1.812 |
| | | | | 0 | 1:90 | beta | 1.812 |
| P0A6E6 | 1AQT | A | 139 | 1 | 91:139 | alpha | 1.4704 |
| | | | | 0\|1 | interface | - | 1.812 |
| P0A763 | 2HUR | C | 143 | 0 | 1:143 | alpha-beta | 1.1556 |
| P77754 | 6BIE | A | 161 | 0 | 1:161 | alpha | 2.0322 |
| P0A7J3 | 6XZ7 | H | 165 | 0 | 1:165 | alpha | 2.0322 |
| P65556 | 2FKB | A | 180 | 0 | 1:180 | alpha-beta | 1.4213 |
| P45748 | 1HRU | A | 190 | 0 | 1:190 | alpha-beta | 1.1556 |
| | | | | 0 | 1:36 / 97:207 | alpha-beta | 1.4213 |
| P60546 | 2ANB | A | 207 | 1 | 37:96 | alpha-beta | 1.9644 |
| | | | | 0\|1 | interface | - | 2.167 |
| P60438 | 6PCR | N | 209 | 0 | 1:209 | beta | 2.1508 |
| P0A6I0 | 1KDO | A | 227 | 0 | 1:227 | alpha-beta | 1.1556 |
| P0A8I5 | 3DXX | A | 239 | 0 | 1:239 | alpha-beta | 1.1556 |
| | | | 281 | 0 | 1:149 | alpha-beta | 1.1556 |
| P31142 | 1URH | A | | 1 | 150:281 | alpha-beta | 1.1556 |
| | | | | 0\|1 | interface | - | 2.5044 |
| P0A6L2 | 1DHP | A | 292 | | 1:292 | alpha-beta | 1.1556 |
| | | | | 0 | 1:58 | alpha | 2.5044 |
| | | | | 1 | 59:160 291:322 | alpha-beta | 1.4213 |
| P0ACP7 | 1QPZ | A | 341 | 2 | 161:290 323:341 | alpha-beta | 1.4213 |
| | | | | 0\|1 | | | 1.2747 |
| | | | | 0\|2 | interface | - | 1.2747 |
| | | | | 1\|2 | | | 1.5679 |
| | | | | 0 | 1:9 218:359 | beta | 1.4732 |
| P0A6B4 | 4WR3 | A | 359 | 1 | 10:217 | alpha-beta | 1.1556 |
| | | | | 0\|1 | interface | - | 2.5044 |
| | | | | 0 | 1:79 182:239 315:367 | alpha-beta | 1.1556 |
| P37747 | 1I8T | A | 367 | 1 | 80:181 240:314 | alpha-beta | 1.1556 |
| | | | | 0\|1 | interface | - | 1.8611 |
| | | | | 0 | 1:204 | alpha-beta | 1.1556 |
| Q46856 | 1OJ7 | D | 387 | 1 | 205:387 | alpha | 1.1954 |
| | | | | 0\|1 | interface | - | 1.2747 |
| | | | | 0 | 336:471 | alpha-beta | 1.4213 |
| | | | | 1 | 1:69 169:335 | alpha-beta | 1.1556 |
| P0AD61 | 4YNG | C | 470 | 2 | 70:168 | beta | 1.4732 |
| | | | | 0\|1 | | - | 1.8611 |
| | | | | 0\|2 | interface | - | 1.2747 |
| | | | | 1\|2 | | - | 1.8611 |
| P21599 | 6K0K | A | 480 | | 1:480 | alpha-beta | 1.1556 |
| | | | | 0 | 1:200 | alpha-beta | 1.1556 |
| P0AES0 | 2IO9 | B | 619 | 1 | 201:619 | alpha-beta | 1.1556 |
| | | | | 0\|1 | interface | - | 1.8611 |

#### Supplementary Table 2.7 | Reference simulations fraction of native contacts ($Q$) statistics

| gene | pdb | chain | $\langle Q \rangle$ | $\sigma$ | 95% C.I. lower | 95% C.I. upper | $\langle Q_{mode} \rangle$ | $Q_{threshold}$ |
|---|---|---|---|---|---|---|---|---|
| P60546 | 2ANB | A | 0.902 | 0.033 | 0.902 | 0.903 | 0.914 | 0.815 |
| P0A8I5 | 3DXX | A | 0.924 | 0.031 | 0.923 | 0.924 | 0.934 | 0.841 |
| P31142 | 1URH | A | 0.901 | 0.031 | 0.901 | 0.901 | 0.903 | 0.811 |
| P0A6L2 | 1DHP | A | 0.938 | 0.021 | 0.938 | 0.938 | 0.941 | 0.880 |
| P0ACP7 | 1QPZ | A | 0.907 | 0.038 | 0.907 | 0.908 | 0.911 | 0.796 |
| P0A6B4 | 4WR3 | A | 0.912 | 0.023 | 0.912 | 0.912 | 0.915 | 0.847 |
| P37747 | 1I8T | A | 0.944 | 0.023 | 0.944 | 0.944 | 0.953 | 0.885 |
| Q46856 | 1OJ7 | D | 0.970 | 0.011 | 0.970 | 0.970 | 0.974 | 0.940 |
| P21599 | 6K0K | A | 0.853 | 0.028 | 0.853 | 0.853 | 0.859 | 0.774 |
| P0AES0 | 2IO9 | B | 0.898 | 0.022 | 0.898 | 0.899 | 0.899 | 0.833 |

*code available https://github.com/obrien-lab-psu/Failure-to-Form_Native_Entanglements

**Supplementary Table 2.8 | Reference simulations fraction of native contacts with a change in non-covalent lasso entanglement ($G$) statistics**

| gene | pdb | chain | $\langle G \rangle$ | $\sigma$ | 95% C.I.. lower | 95% C.I. upper | $\langle G_{mode} \rangle$ | $G_{threshold}$ |
|---|---|---|---|---|---|---|---|---|
| P60546 | 2ANB | A | 0.0039 | 0.0026 | 0.0039 | 0.0039 | 0.0039 | 0.0118 |
| P0A8I5 | 3DXX | A | 0.0002 | 0.0011 | 0.0002 | 0.0002 | 0.0000 | 0.0032 |
| P31142 | 1URH | A | 0.0005 | 0.0010 | 0.0005 | 0.0005 | 0.0003 | 0.0033 |
| P0A6L2 | 1DHP | A | 0.0012 | 0.0010 | 0.0012 | 0.0012 | 0.0008 | 0.0038 |
| P0ACP7 | 1QPZ | A | 0.0003 | 0.0007 | 0.0003 | 0.0003 | 0.0000 | 0.0021 |
| P0A6B4 | 4WR3 | A | 0.0012 | 0.0022 | 0.0011 | 0.0012 | 0.0000 | 0.0065 |
| P37747 | 1I8T | A | 0.0006 | 0.0010 | 0.0006 | 0.0006 | 0.0006 | 0.0035 |
| Q46856 | 1OJ7 | D | 0.0008 | 0.0009 | 0.0008 | 0.0008 | 0.0006 | 0.0034 |
| P21599 | 6K0K | A | 0.0028 | 0.0031 | 0.0027 | 0.0028 | 0.0008 | 0.0102 |
| P0AES0 | 2IO9 | B | 0.0009 | 0.0012 | 0.0009 | 0.0009 | 0.0007 | 0.0042 |

*code available https://github.com/obrien-lab-psu/Failure-to-Form_Native_Entanglements

#### Supplementary Table 2.9 | Reference simulations mirror metric ($K$)

| gene | pdb | chain | $\langle K \rangle$ | $\sigma$ | 95% C.I. lower | 95% C.I. upper | $\langle K_{mode} \rangle$ |
|---|---|---|---|---|---|---|---|
| P60546 | 2ANB | A | 0.837 | 0.074 | 0.837 | 0.838 | 0.852 |
| P0A8I5 | 3DXX | A | 0.888 | 0.073 | 0.887 | 0.888 | 0.924 |
| P31142 | 1URH | A | 0.858 | 0.056 | 0.858 | 0.859 | 0.882 |
| P0A6L2 | 1DHP | A | 0.949 | 0.035 | 0.949 | 0.949 | 0.973 |
| P0ACP7 | 1QPZ | A | 0.875 | 0.048 | 0.875 | 0.875 | 0.892 |
| P0A6B4 | 4WR3 | A | 0.852 | 0.042 | 0.852 | 0.852 | 0.851 |
| P37747 | 1I8T | A | 0.927 | 0.037 | 0.926 | 0.927 | 0.936 |
| Q46856 | 1OJ7 | D | 0.938 | 0.039 | 0.938 | 0.939 | 0.959 |
| P21599 | 6K0K | A | 0.875 | 0.044 | 0.874 | 0.875 | 0.876 |
| P0AES0 | 2IO9 | B | 0.862 | 0.033 | 0.862 | 0.863 | 0.865 |

*code available https://github.com/obrien-lab-psu/Failure-to-Form_Native_Entanglements

#### Supplementary Table 2.10 | Optimal set of non-covalent lasso entanglement properties to classify proteins based on essentiality in the absence of chaperones

| | | Essential proteins | | | | Non-essential proteins | | | | | | | |
|---|---|---|---|---|---|---|---|---|---|---|---|---|---|
| **Feature** | **ID** | **Mean** | **C.I.** | **Median** | **C.I.** | **Mean** | **C.I.** | **Median** | **C.I.** | **MWU p-values (FDR)** | **Perm. p-values (FDR)** | **Mean relative difference** | **Median relative difference** |
| $N_{\mathrm{Threads}}$ | 1 | 1.19 | (1.11, 1.32) | 1 | (1, 3) | 1.21 | (1.14, 1.30) | 1 | (1, 3) | 1 | 1 | -1.1 | 0.0 |
| $g_{\mathrm{c}}$ | 2 | 0.78 | (0.76, 0.83) | 0.75 | (0.60, 1.49) | 0.85 | (0.82, 0.88) | 0.79 | (0.61, 1.68) | 0.030 | 0.120 | -7.7 | -6.2 |
| $C_{\mathrm{Threads}}$ | 3 | 1.14 | (1.07, 1.29) | 1.00 | (1.00, 3.00) | 1.15 | (1.10, 1.22) | 1.00 | (1.00, 3.00) | 1 | 1 | -0.3 | 0.0 |
| $N_{\mathrm{Loop}}$ | 4 | 98.2 | (87.0, 111.9) | 64.00 | (16.0, 407.0) | 99.03 | (92.2, 107.1) | 63.00 | (17, 328) | 1 | 1 | -0.9 | 1.6 |
| $N_{\mathrm{Loop-cont}}$ | 7 | 153.3 | (140.7, 168.7) | 115.00 | (45.0, 495.0) | 158.21 | (149.9, 167.5) | 122.00 | (48, 463) | 1 | 1 | -3.1 | -5.9 |
| $N_{\mathrm{Cross-cont}}$ | 8 | 39.8 | (38.1, 42.0) | 36.00 | (27.0, 88.0) | 41.56 | (40.2, 43.2) | 36.00 | (27.0, 93.0) | 0.421 | 1 | -4.3 | 0.0 |
| $f_{\mathrm{entR}}$ | 9 | 0.18 | (0.17, 0.19) | 0.15 | (0.04, 0.42) | 0.20 | (0.19, 0.21) | 0.18 | (0.07, 0.44) | 0.003 | 0.108 | -10.4 | -18.4 |
| $\mathrm{MIN}[d_{\mathrm{P,N}}]$ | 10 | 0.30 | (0.27, 0.34) | 0.26 | (0.02, 0.77) | 0.38 | (0.35, 0.40) | 0.38 | (0.02, 0.82) | 0.014 | 0.031 | -20.9 | -36.6 |
| $\mathrm{MIN}[d_{\mathrm{T,N}}]$ | 11 | 0.71 | (0.66, 0.75) | 0.82 | (0.10, 0.98) | 0.75 | (0.73, 0.78) | 0.85 | (0.14, 0.98) | 0.661 | 1.034 | -5.9 | -3.6 |
| $\mathrm{MIN}[d_{\mathrm{S,C}}]$ | 12 | 181.1 | (151.4, 217.1) | 124.00 | (8.0, 696.0) | 173.34 | (156.9, 191.5) | 131.00 | (9.0, 613.0) | 1 | 1 | 4.3 | -5.4 |
| $\mathrm{MIN}[d_{\mathrm{S,C}}]$ | 15 | 224.1 | (188.0, 267.7) | 128.00 | (14.0, 750.0) | 147.01 | (131.8, 165.2) | 103.00 | (13.0, 497) | 0.015 | 0.002 | 41.5 | 21.6 |
| $RCO$ | 17 | 0.28 | (0.25, 0.30) | 0.22 | (0.02, 0.74) | 0.30 | (0.29, 0.32) | 0.25 | (0.06, 0.80) | 0.108 | 0.946 | -9.8 | -13.7 |

* For the tables of optimal features in the presence of chaperones see Supplementary information file 6

### 3. Supplemental Figures

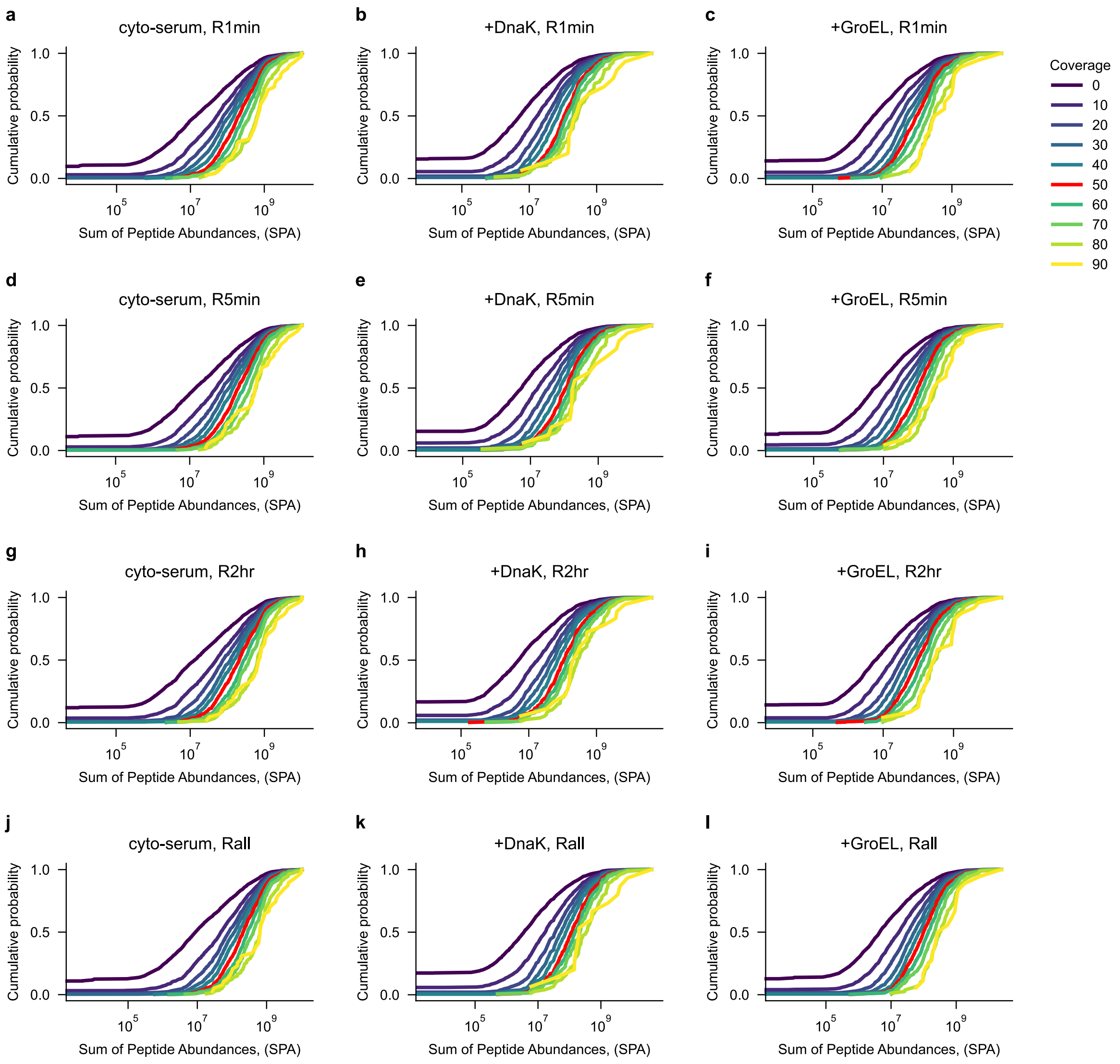

**Supplementary Fig. 1 | Sum of the peptide abundance (SPA) cumulative probability in the untreated sample dataset as a function of primary structure coverage. a,** each protein observed in the proteome wide limited proteolysis dataset has a pool of peptides that each have a quantified abundance. We use the sum of the peptide abundance for a protein as a proxy for its natural abundance in the cell. For the untreated sample we rank order the proteins by their SPA and then use it as a convenient scale to test our conclusions robustness to protein abundance changes in the cell. **a-l**, cumulative density functions of the untreated samples SPA at various buffer conditions and refolding time points. Within each condition and time point we vary how strict the coverage of the primary structure threshold is required to be a viable candidate in our dataset. All work presented in the main text requires at least 50% of the primary sequence be resolved.

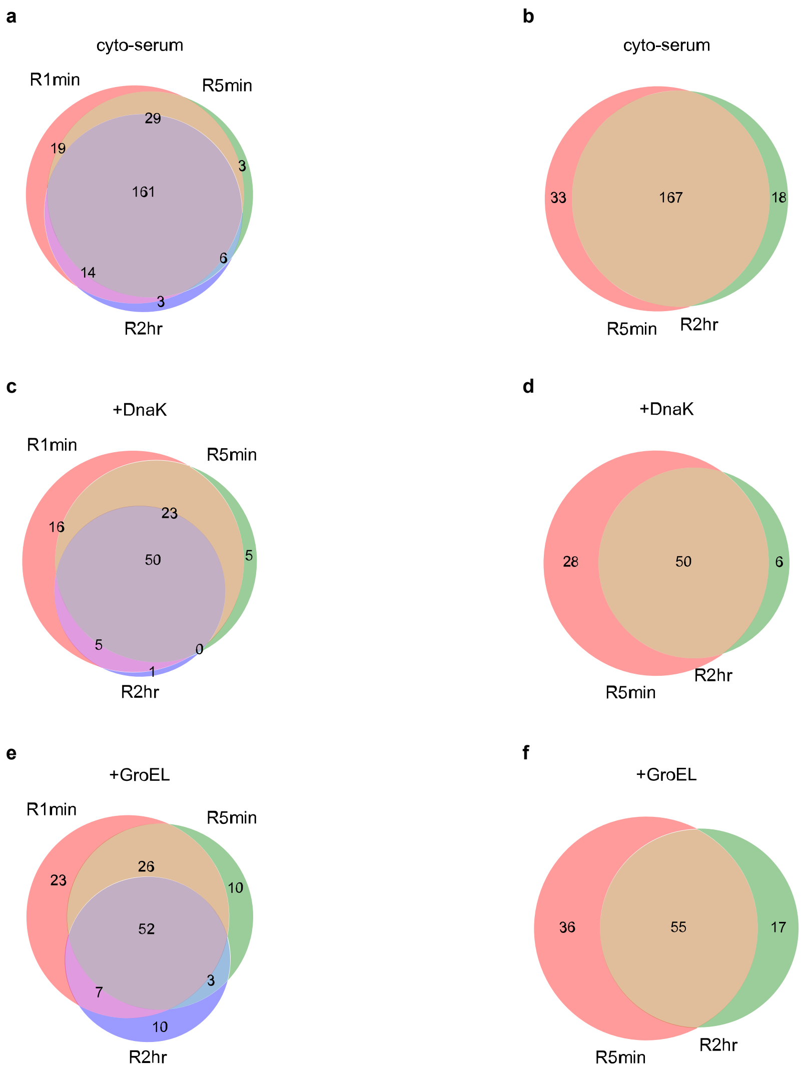

**Supplementary Fig. 2 | Overlap of proteins containing non-covalent lasso entanglements with significant changes in structure observed at various refolding times and buffer conditions a, c, e,** The overlap of proteins containing native NCLEs observed at 1 min, 5 min, and 2 h refolding times in the absence of chaperones and the presence of +DnaK and +GroEL respectively. **b, d, f,** the same as in **a, c,** and **e,** but considering only the later two refolding times of 5 min and 2 h. All overlaps are significant beyond random chance with p-values <= 0.0001.

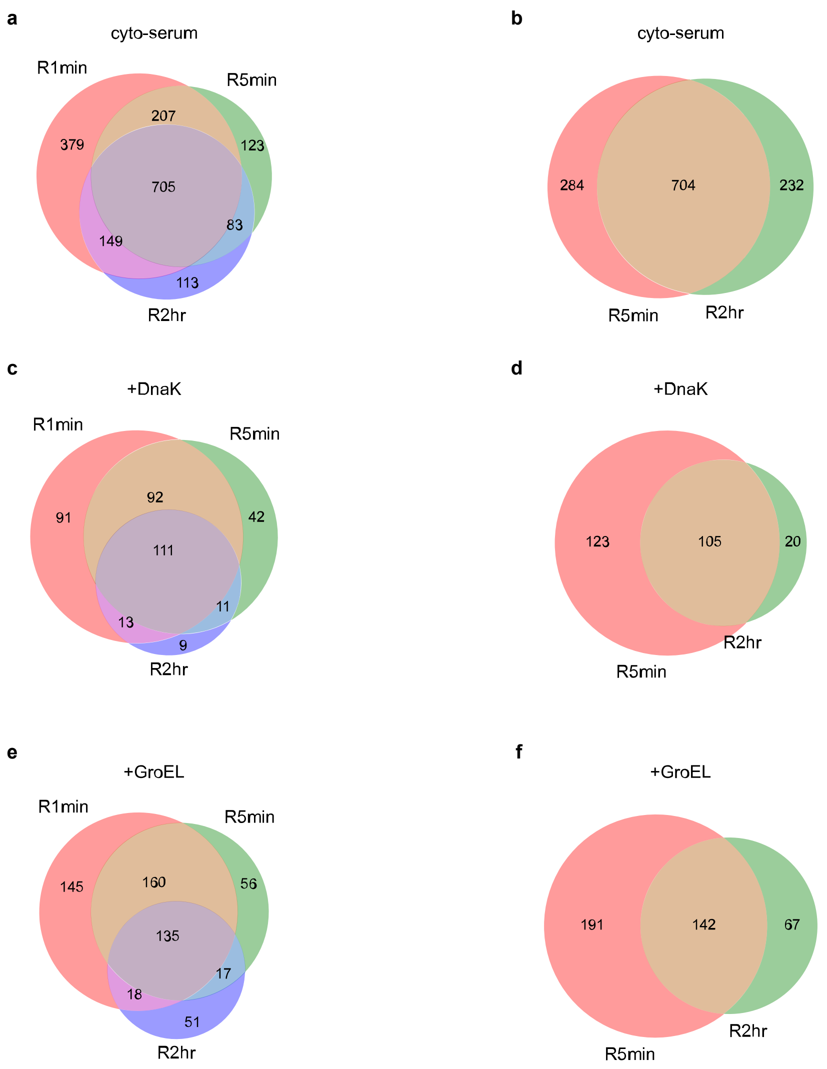

**Supplementary Fig. 3 | Overlap of significant PK cut-sites in proteins observed at various refolding times and buffer conditions a,** The overlap of significant proteinase-K cut-sites in proteins containing native NCLEs observed at 1 min, 5 min, and 2 h refolding times in the absence of chaperones and the presence of +DnaK and +GroEL respectively. Overlap was determined if the cut-sites were within +/- 3 residues of each other along the primary structure. **b, d, f,** the same as in **a, c,** and **e,** but considering only the later two refolding times of 5 min and 2 h. All overlaps are significant beyond random chance with p-values <= 0.0001 with the exception of **f** where the p-value = 0.06.

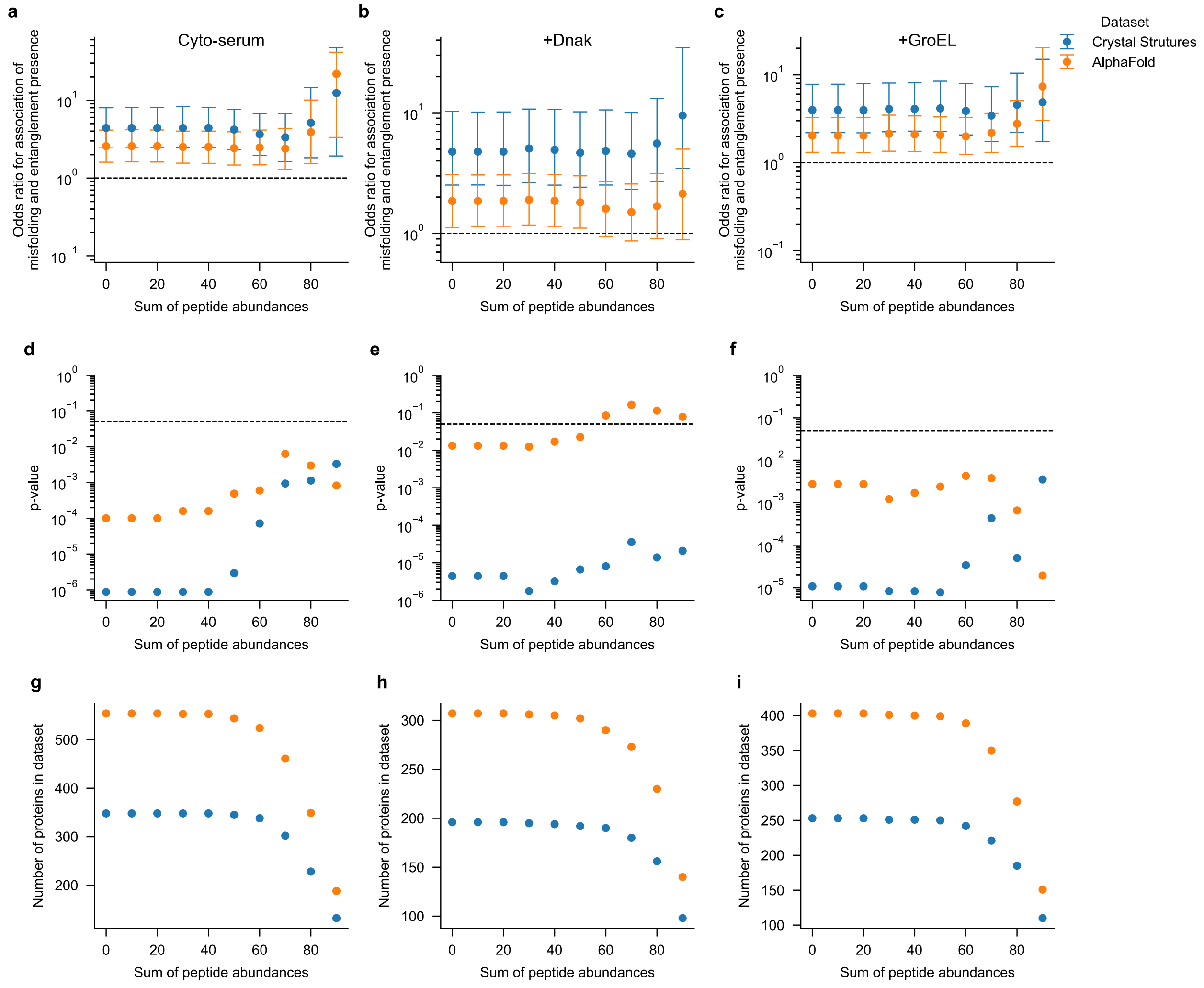

**Supplementary Fig. 4 | Robustness of odds ratio for the association of misfolding with the presence of native non-covalent lasso entanglements as a function of SPA.** we repeat the maximum likelihood fit on subsets of the proteins that meet or exceed a certain SPA threshold set by the natural percentiles of the SPA distribution in the untreated samples. **a,** odds ratios resulting from the maximum likelihood fit of a logistic regression model of the log odds of a protein having at-least 1 significant cut-site as a function of the protein length and presence of a native NCLE. **b,** same as in **a** but for the buffer containing the chaperone DnaK. **c,** same as in **b** but with GroEL. **d, e, f,** are the respective p-values derived from the model fit using the Wald test. **g, h, j** are the respective sample sizes in each percentile assessed.

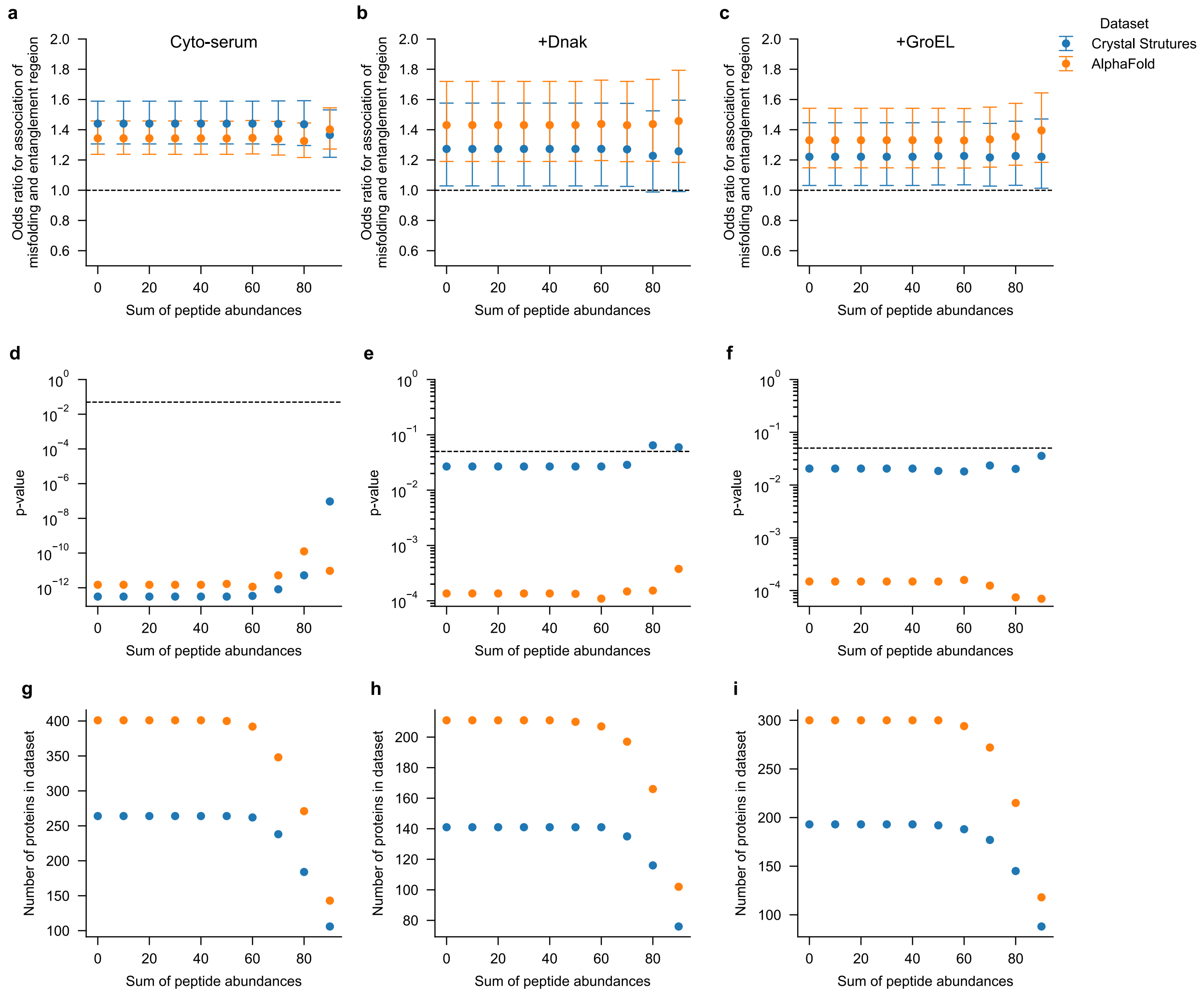

**Supplementary Fig. 5 | Robustness of odds ratio for the association of misfolding and non-covalent lasso entanglement region as a function of SPA.** we repeat the maximum likelihood fit on subsets of the proteins that meet or exceed a certain SPA threshold set by the natural percentiles of the SPA distribution in the untreated samples. **a,** odds ratios resulting from the maximum likelihood fit of a logistic regression model of the log odds of observing a significant cut-site as a function of that site's amino acid identity and whether it was located in the natively entangled region or not. **b,** same as in **a** but for the buffer containing the chaperone DnaK. **c,** same as in **b** but with GroEL. **d, e, f,** are the respective p-values derived from the model fit using the Wald test. **g, h, j** are the respective sample sizes in each percentile assessed.

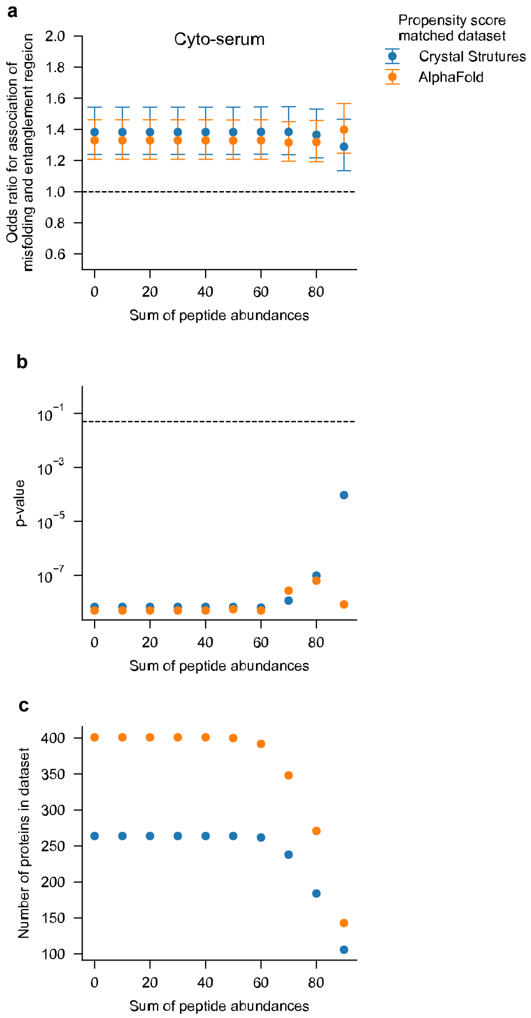

**Supplementary Fig. 6 | Robustness of odds ratio for the association of misfolding and non-covalent lasso entanglement region as a function of SPA after controlling for solvent accessibility.** Each residue in the entangled region of proteins in our dataset was matched to one from the non-entangled region using the solvent accessible surface area and the propensity score matching method in R. We repeat the maximum likelihood fit on subsets of the proteins that meet or exceed a certain SPA threshold set by the natural percentiles of the SPA distribution in the untreated samples. **a,** odds ratios resulting from the maximum likelihood fit of a logistic regression model of the log odds of observing a significant cut-site as a function of that site's amino acid identity and whether it was located in the natively entangled region or not. **b,** the p-values derived from the model fit using the Wald test. **c,** are the sample sizes in each percentile assessed.

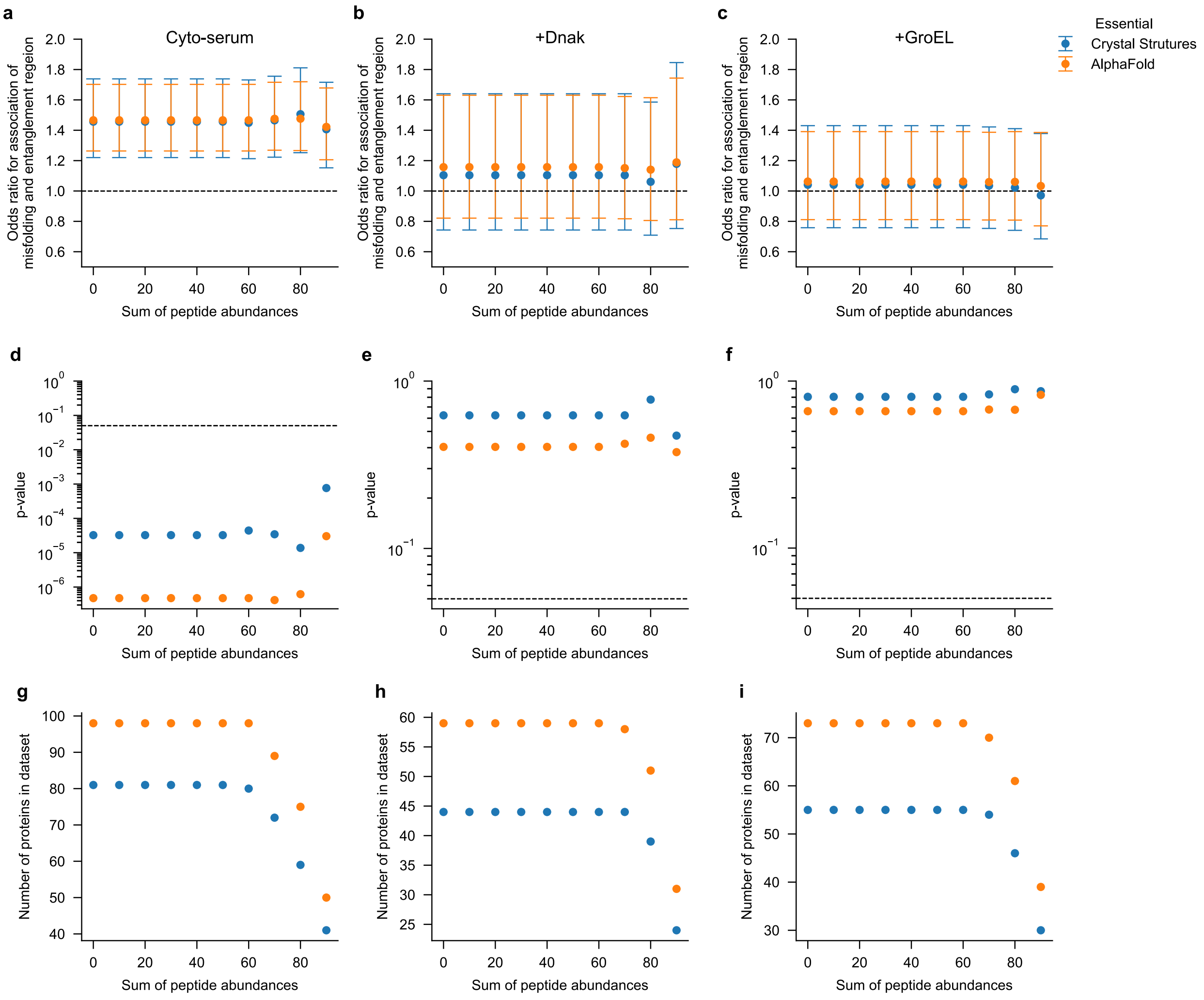

**Supplementary Fig. 7 | Robustness of odds ratio for the association of misfolding and non-covalent lasso entanglement region as a function of SPA for essential proteins containing native non-covalent lasso entanglements.** We repeat the maximum likelihood fit on subsets of the essential proteins that meet or exceed a certain SPA threshold set by the natural percentiles of the SPA distribution in the untreated samples **a,** odds ratios resulting from the maximum likelihood fit of a logistic regression model of the log odds of observing a significant cut-site as a function of that site's amino acid identity and whether it was located in the natively entangled region or not. **b,** same as in **a** but for the buffer containing the chaperone DnaK. **c,** same as in **b** but with GroEL. **d, e, f,** are the respective p-values derived from the model fit using the Wald test. **g, h, j** are the respective sample sizes in each percentile assessed.

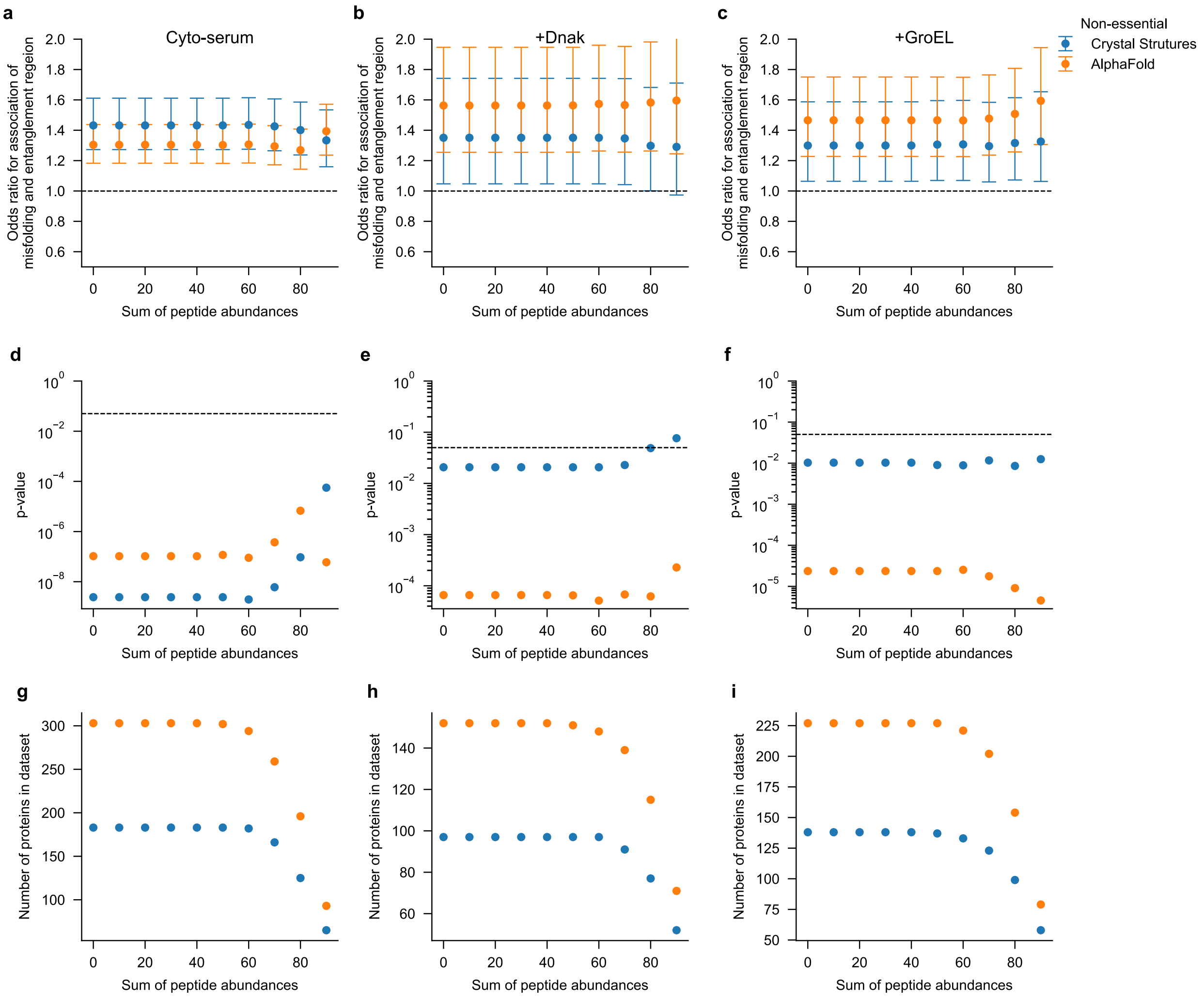

**Supplementary Fig. 8 | Robustness of odds ratio for the association of misfolding and non-covalent lasso entanglement region as a function of SPA for non-essential proteins containing native non-covalent lasso entanglements. a,** we repeat the maximum likelihood fit on subsets of the non-essential proteins that meet or exceed a certain SPA threshold set by the natural percentiles of the SPA distribution in the untreated samples **a,** odds ratios resulting from the maximum likelihood fit of a logistic regression model of the log odds of observing a significant cut-site as a function of that site's amino acid identity and whether it was located in the natively entangled region or not. **b,** same as in **a** but for the buffer containing the chaperone DnaK. **c,** same as in **b** but with GroEL. **d, e, f,** are the respective p-values derived from the model fit using the Wald test. **g, h, j** are the respective sample sizes in each percentile assessed.

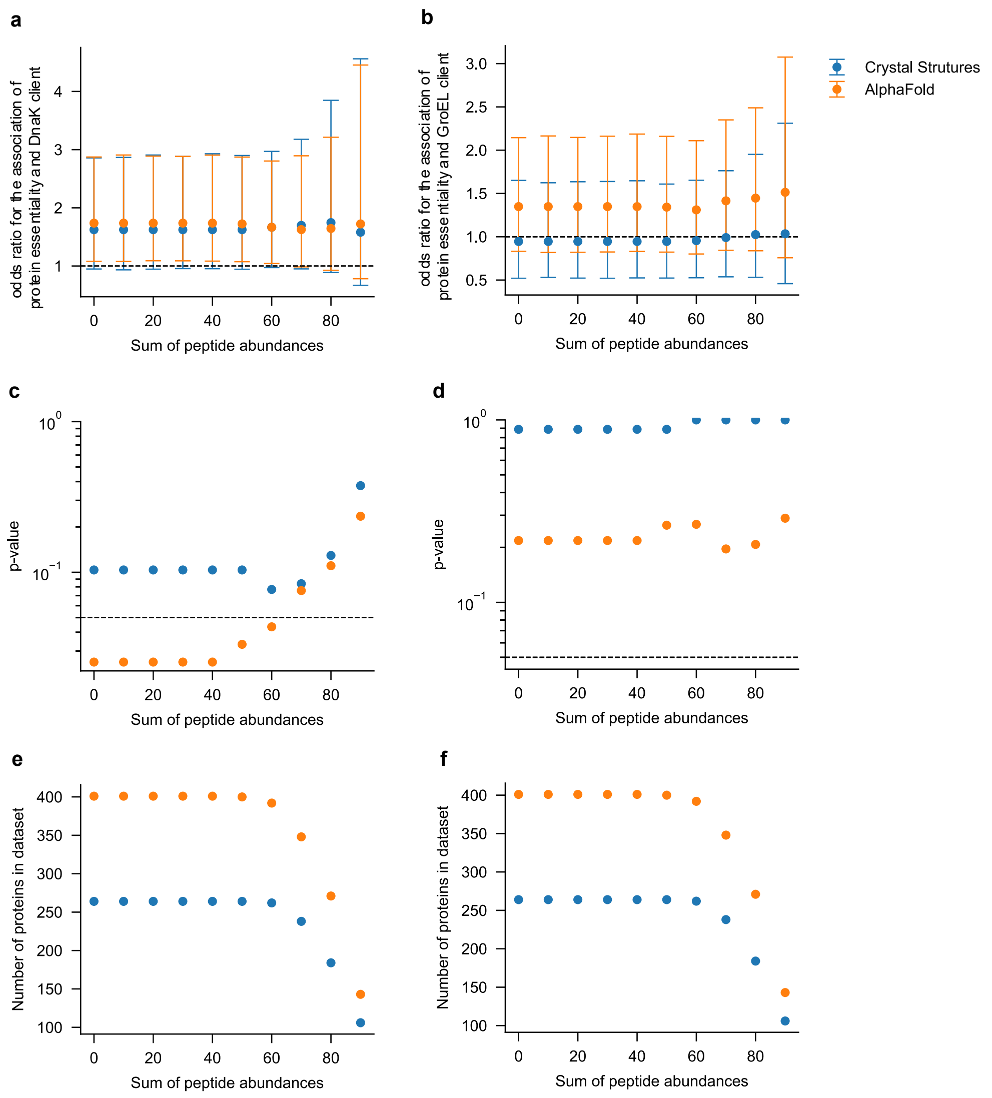

**Supplementary Fig. 9 | Robustness of odds ratio for the association of proteins containing native non-covalent lasso entanglement essentiality and being a known client of DnaK. a,** we test for significant association of protein essentiality and being a known DnaK or GroEL client with the Fisher Exact test applied on subsets of the proteins that meet or exceed a certain SPA threshold set by the natural percentiles of the SPA distribution in the untreated samples **a, b,** odds ratios resulting from a contingency table defined by protein essentiality and whether the protein is a known client of DnaK or GroEL respectfully. The odds ratio is defined $\mathrm{OR} = \frac{\mathrm{P}_{11}\mathrm{P}_{00}}{\mathrm{P}_{10}\mathrm{P}_{01}}$, where $\mathrm{P}_{11}$ is the probability of a protein both containing a native NCLE and being non-refoldable, $\mathrm{P}_{00}$ is the probability of a protein not containing a native NCLE and being refoldable, $\mathrm{P}_{10}$ is the probability of a protein containing a native NCLE and being refoldable, $\mathrm{P}_{01}$ is the probability of a protein not containing a native NCLE and being non-refoldable. We estimate the OR from fitting a binomial logistic regression model. **d, e, f,** are the respective p-values derived from the model fit using the Wald test. **g, h, j** are the respective sample sizes in each percentile assessed.

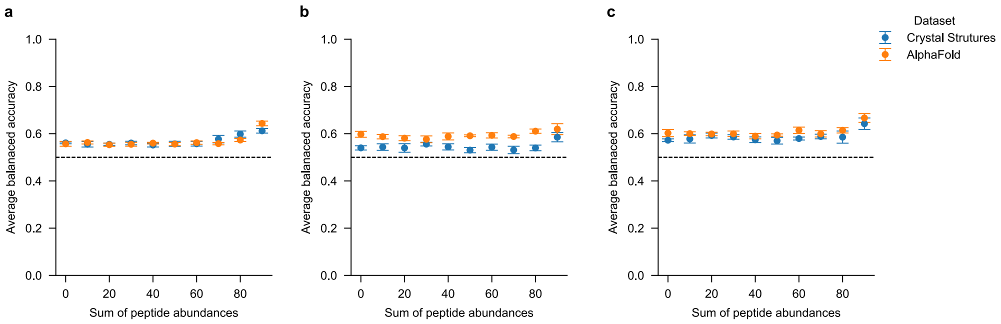

**Supplementary Fig. 10 | Robustness of model predicting protein essentiality as a function of non-covalent lasso entanglement topology.** The least absolute shrinkage and selection operator (L.A.S.S.O) regression analysis was applied on subsets of the proteins that meet or exceed a certain SPA threshold set by the natural percentiles of the SPA distribution in the untreated samples. The maximal balanced accuracies are reported as a measure of the power of NCLE structural features to discriminate between essential and non-essential proteins. **a,** in the absence of chaperones, **b,** in the presence of DnaK, and **c,** in the presence of GroEL.

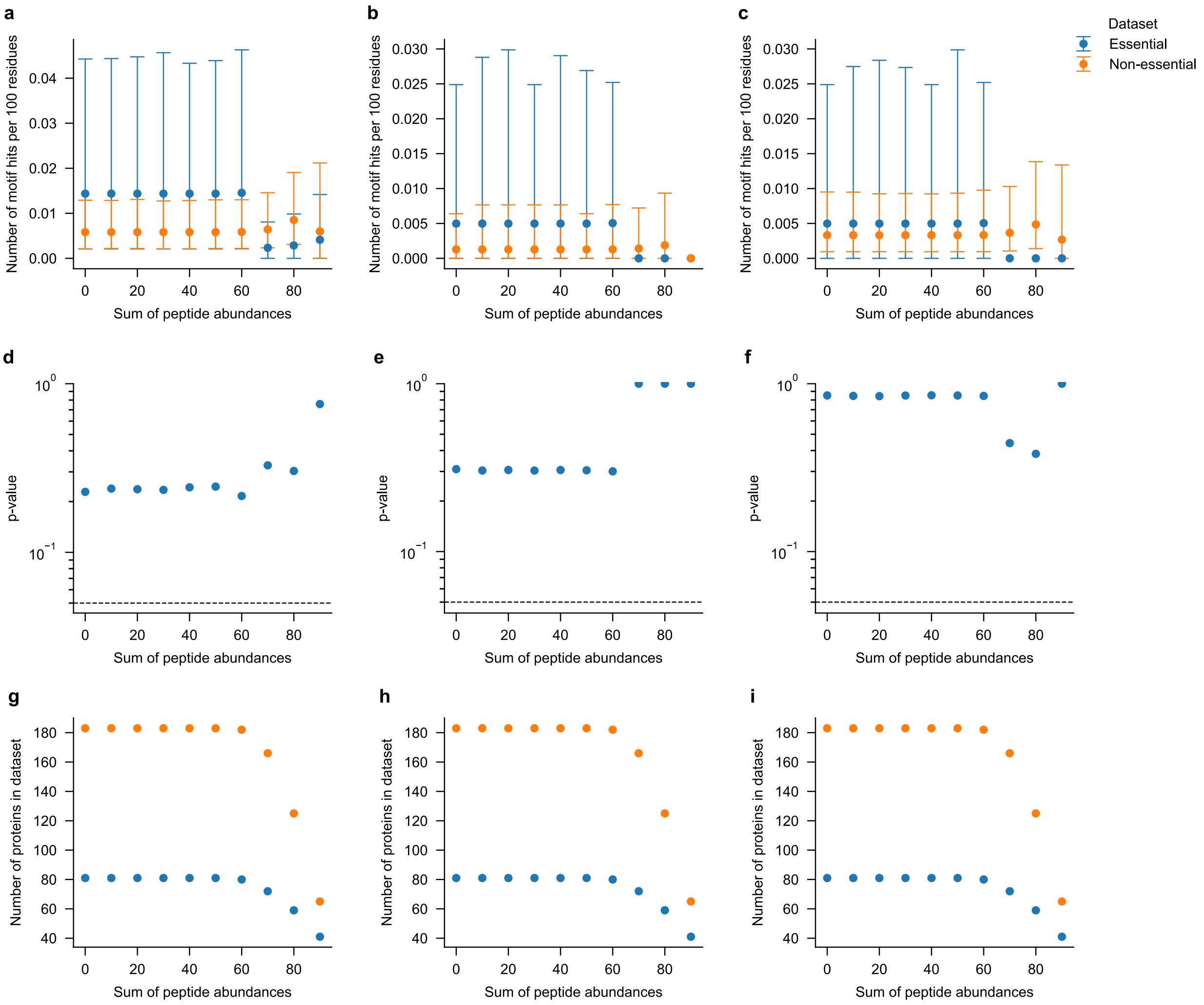

**Supplementary Fig. 11 | Robustness of sequence scan for DnaK binding motifs a,** we calculate the number of identified and predicted DnaK binding motifs[12,13] present along the canonical protein sequences per every 100 residues for subsets of the proteins that meet or exceed a certain SPA threshold set by the natural percentiles of the SPA distribution in the untreated samples. The allowed motifs are defined in supplementary table S2.1 and the motif presented here is the (Schymkowitz motif) as it includes information both from experiments and homology modeling. **b,** same as in **a** but for only the threading regions of the native NCLEs of each protein. **c,** same as in **a** but for only the loop regions of proteins. **d,** the p-value for a permutation test comparing if the motif hit rate is significantly different between essential and non-essential proteins. **e, f,** same as in f but for the threading region and loop closing regions respectively. **g**, the sample size at each SPA threshold. **h,i,** same as in **g** but for the threading region and loop closing regions respectively.

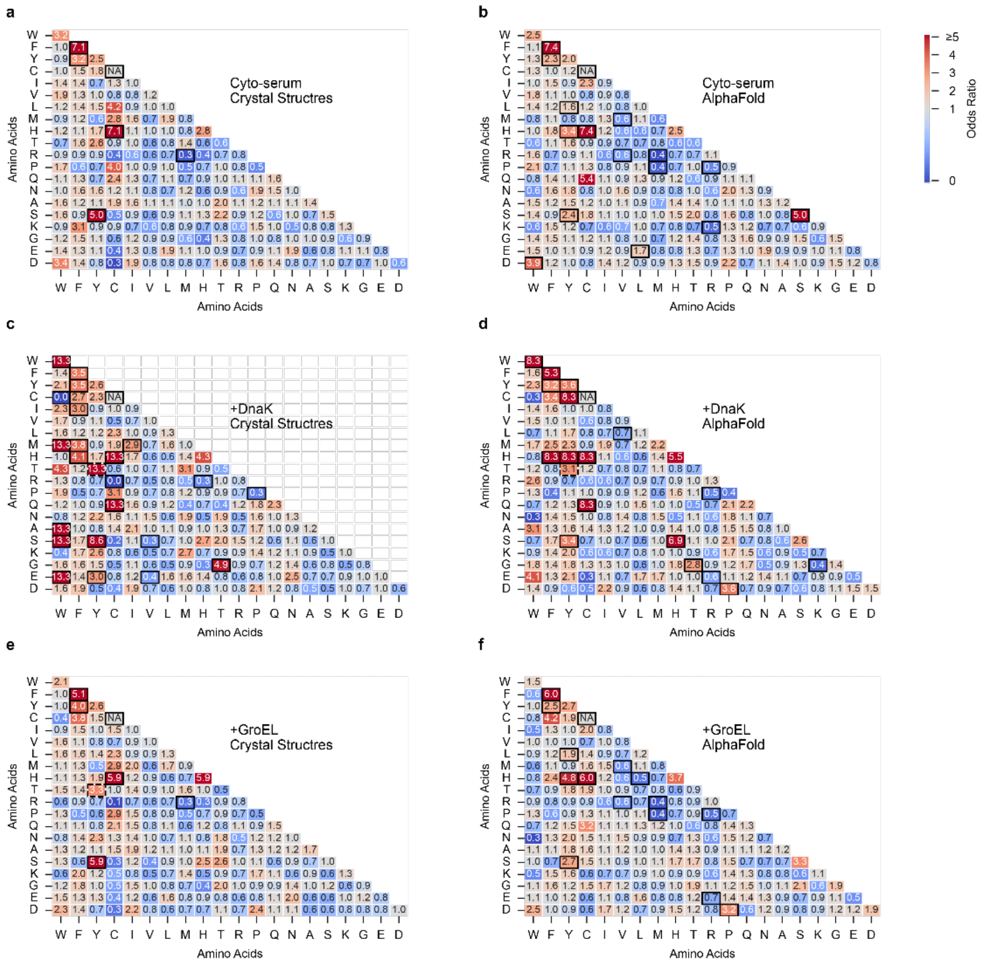

**Supplementary Fig. 12 | Loop forming native contact identity enrichment in non-essential proteins.** test for significant enrichment of a particular pair of amino acids closing the base of the native NCLE loop in non-essential proteins relative to essential proteins on subsets of the proteins that meet or exceed a certain SPA threshold set by the natural percentiles of the SPA distribution in the untreated samples **a-f,** for each pair of loop closing contacts we calculate the odds ratios resulting from a contingency table defined by parent protein essentiality and whether the contact is made up of particular amino acid pairs. The odds ratio is defined $\mathrm{OR} = \frac{\mathrm{P}_{11}\mathrm{P}_{00}}{\mathrm{P}_{10}\mathrm{P}_{01}}$, where $\mathrm{P}_{11}$ is the probability of the contact originates from a non-essential parent protein and being of the type $(i, j)$, $\mathrm{P}_{00}$ is the probability of the contact originating from an essential parent protein and not being of the type $(i, j)$, $\mathrm{P}_{10}$ is the probability of the contact originates from a non-essential parent protein and of not being of the type $(i, j)$, $\mathrm{P}_{01}$ is the probability of the contact originates from an essential parent protein and being of the type $(i, j)$. We calculate p-values using classic permutation ($n$ = 100,000). Black boxed amino acids indicate those significantly enriched beyond random chance.

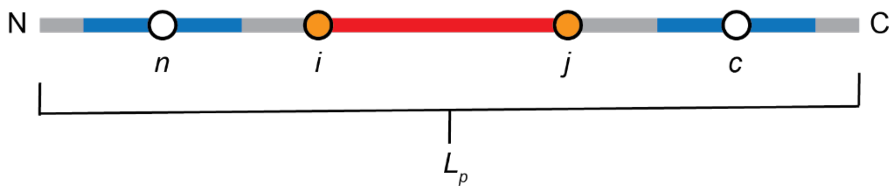

**Supplementary Fig. 13 | Schematic of a native non-covalent lasso entanglement defining variables used in NCLE structural parameters.** Schematic to define key parameters used to quantify topological features of a native NCLE. The loop (red) closing residues are show as orange spheres labeled $i$ and $j$. The segments of the backbone that thread through the loop are colored in blue and the residues that cross the plane of the loop are shown as white spheres and labeled $n$ and $c$ for their respective termini. $L_p$ is the length of the protein chain.

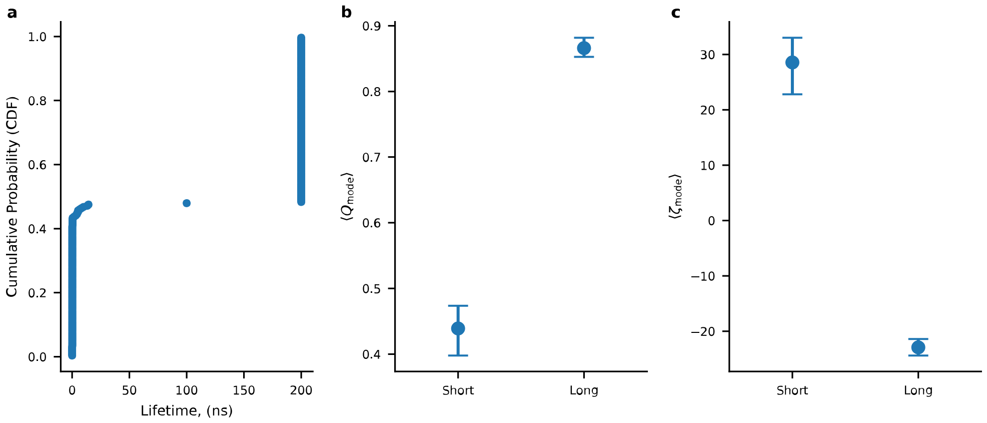

**Supplementary Fig. 14 | Estimated lifetime, native structure, and native hydrophobic exposure of simulated misfolded states of proteins that contained native NCLEs. a,** The cumulative density function of the average misfolded state lifetimes across all temperature quenching trajectories. Trajectories are then grouped into short and long lived states depending on if they have an average life time greater than 100 ns. b, the average $Q_{\mathrm{mode}}$ derived from trajectory sampling via a sliding window so 15 ns. c, the average $\zeta_{\mathrm{mode}}$ derived from trajectory sampling via a sliding window so 15 ns.

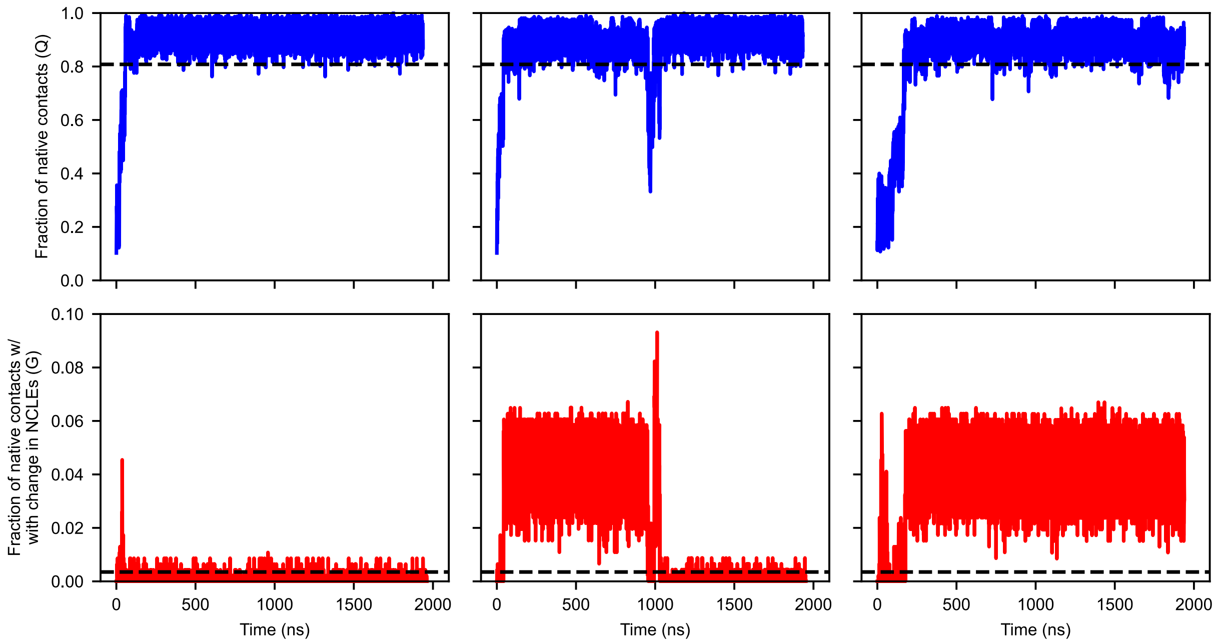

**Supplementary Fig. 15 | Select trajectories fraction of native contacts (Q) and fraction of native contacts with a change in NCLE status (G) time series**. **(top)** Q for the last 200 ns of each temperature quenching trajectory for P0A8I5 with the folded threshold of 0.841 (black dashed lines) **(bottom)** G for the last 200 ns of each temperature quenching trajectory for P0A8I5 with the folded threshold of 0.0032 (black dashed lines) **(left)** – example of a folded and highly native trajectory, **(middle)** example of a misfolded and highly native trajectory, **(right)** example of a misfolded non-native trajectory.

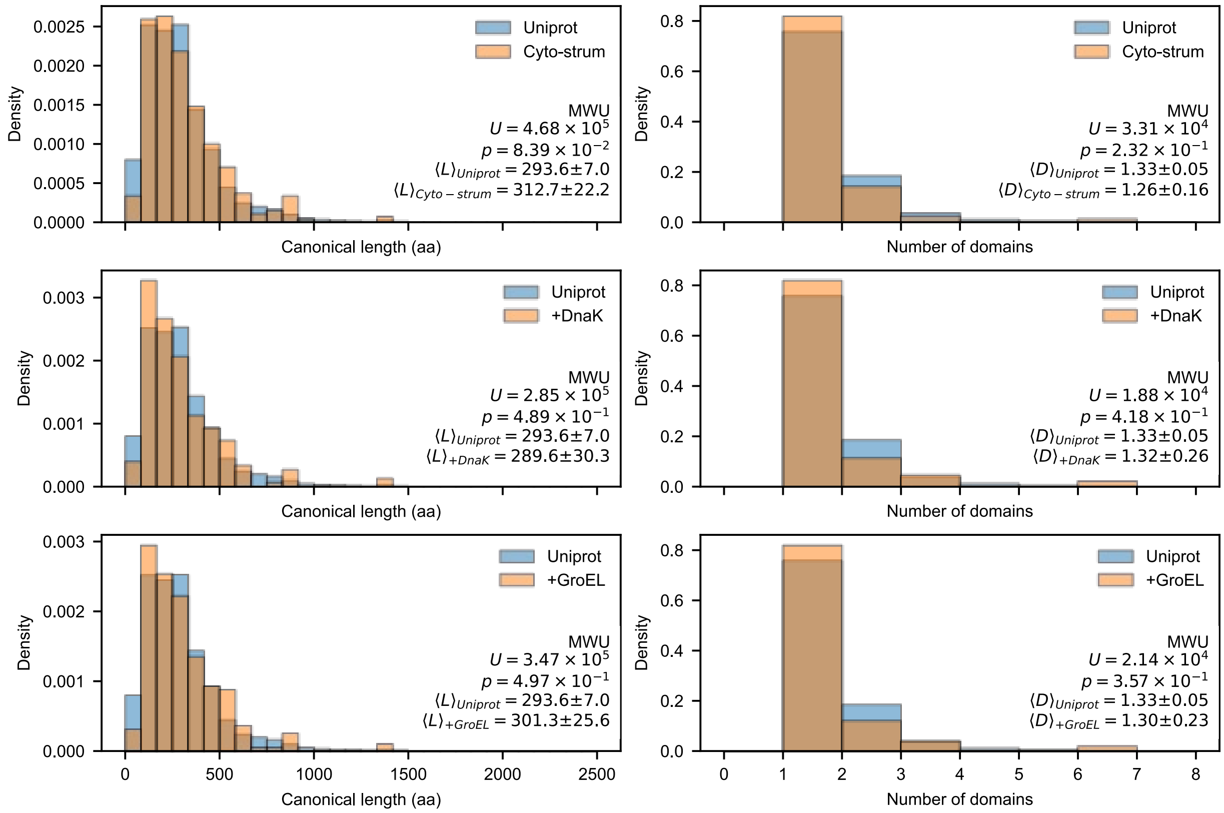

**Supplementary Fig. 16 | Protein length and domain count representative of whole *E. coli* proteome (experimental structures)**. **(left)** protein size distributions of each set of proteins observed in the LiP-MS experiments (Cyto-serum only, +DnaK, and +GroEL) with high quality PDB structures (orange) compared with the whole reviewed *E. coli* cytosolic protoeme from Uniprot (blue, $n = 3{,}069$). **(right)** domain count distributions of each set of proteins observed in the LiP-MS experiments (Cyto-serum only, +DnaK, and +GroEL) with high quality PDB structures (orange) compared with the whole reviewed *E. coli* cytosolic proteome from Uniprot (blue, $n = 809$). Mann Whitney U-test was used to determine if the two distributions differed significantly.

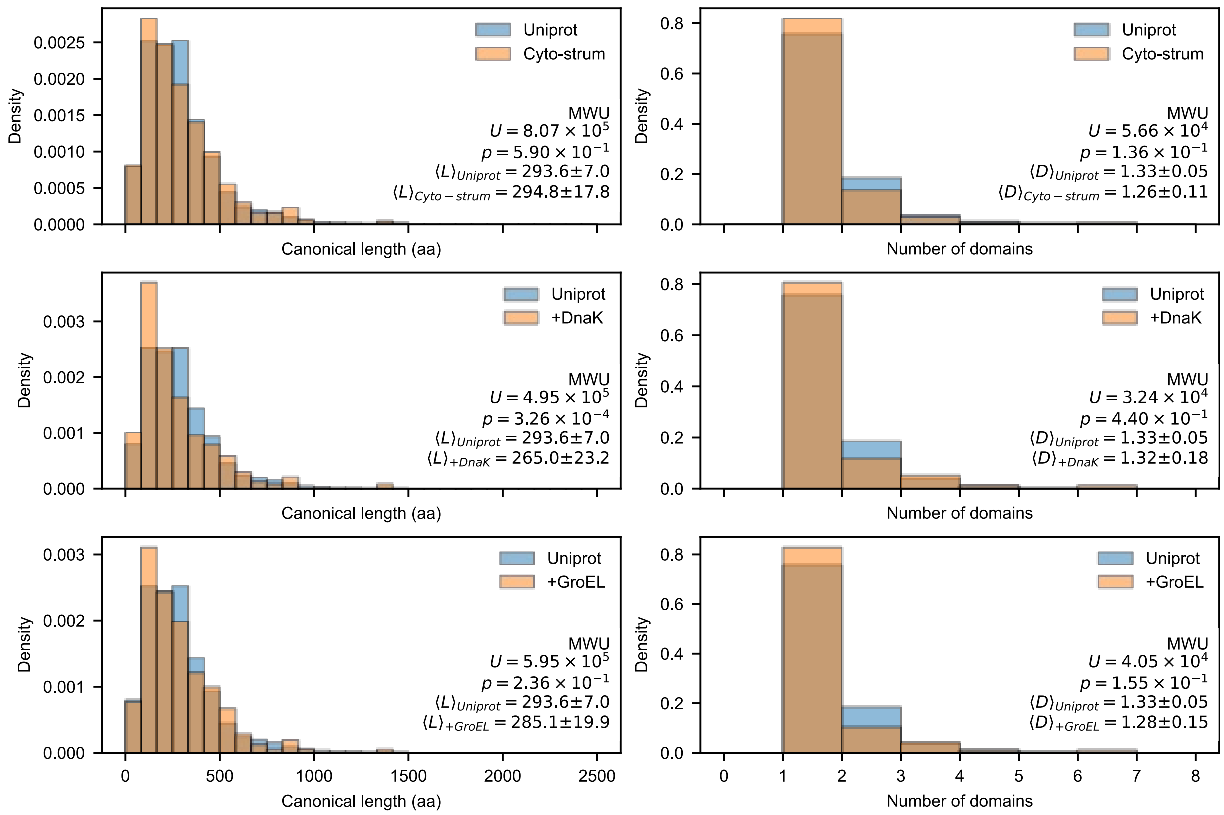

**Supplementary Fig. 17 | Protein length and domain count representative of whole *E. coli* proteome (Alpha Fold structures)**. **(left)** protein size distributions of each set of proteins observed in the LiP-MS experiments (Cyto-serum only, +DnaK, and +GroEL) with high quality AF structures (orange) compared with the whole reviewed *E. coli* cytosolic protoeme from Uniprot (blue, $n = 3{,}069$). **(right)** domain count distributions of each set of proteins observed in the LiP-MS experiments (Cyto-serum only, +DnaK, and +GroEL) with high quality AF structures (orange) compared with the whole reviewed *E. coli* cytosolic protoeme from Uniprot (blue, $n = 809$). Mann Whitney U-test was used to determine if the two distributions differed significantly.
